# Robust CRISPR/dCas13 RNA blockers specifically perturb miRNA-target interactions and rescue type 1 myotonic dystrophy pathology

**DOI:** 10.1101/2024.09.16.612263

**Authors:** Muhammad Hanifi, Perihan Seda Ates-Kalkan, Sean Wen, Mathieu Fischer, Amanda Kroesen, Zulin Yu, Matthew Wood, Supat Thongjuea, Adam Mead, Tudor Alexandru Fulga, Carlo Rinaldi, Tatjana Sauka-Spengler

## Abstract

While RNA-targeting strategies are powerful tools for disease therapy, challenges, including low target engagement and off-target collateral effects, currently limit their efficacy. Here, we report the engineering and optimisation of a CRISPR/dCas13 RNA steric blocker (CRISPR-Lock) that prevents mRNA translation, shields mRNAs from miRNA-mediated silencing, and blocks RNA-protein interactions. By tuning the spatial resolution and mismatch tolerance of CRISPR-Lock, we develop a high-resolution perturbation approach that employs genetically encoded CRISPR-Lock as a miRNA target protector. This system enables precise spatiotemporal control of miRNA:mRNA interactions, offering broader applicability compared to phosphorodiamidate morpholino (PMO) target protectors. Moreover, we demonstrate the potential therapeutic application of CRISPR-Lock for blocking pathological RNA-protein interactions in type 1 myotonic dystrophy (DM1). Optimising CRISPR-Lock to target expanded repeat RNAs corrects approximately 85% of clinically relevant splicing biomarkers in patient-derived myotubes and significantly out-performs third-generation PMO antisense oligonucleotides. Finally, by delivering a miniaturised AAV-encoded CRISPR-Lock system into an established DM1 mouse model, we demonstrate the dose-dependent correction of intranuclear foci and splicing dysregulation, underscoring the potential therapeutic application of this technology.

## 1 Introduction

As central mediators of genetic information flow, RNAs have long been recognised as a potential therapeutic target for a broad spectrum of diseases^1–4^. Coding and non-coding RNAs assume diverse roles inside cells, such as providing templates for protein translation, tuning gene expression^5–7^, modulating protein activities^8,9^, and acting as molecular scaffolds^10,11^. These broad roles explain why RNA dysregulation has been implicated in many diseases, including malignancies^12–14^, metabolic^8,9^, cardiovascular^15–17^, and neurological disorders^18–20^. Tapping into the potential of RNAs as therapeutic targets, many RNA-targeting modalities using small molecules^2,21^, antisense oligonucleotides (ASOs)^22,23^, aptamers^24,25^, siRNAs^26,27^, miRNAs^28,29^, and more recently, synthetic mRNAs^30,31^ have been developed. In the last decade, a new cohort of RNA therapeutics has received FDA and EMA approvals for metabolic diseases, rare genetic diseases, and COVID-19 infection^3^.

Despite these rapid developments, the promise of RNA medicine for the modulation of any cellular RNA remains hindered by several important obstacles. Oligonucleotide-based RNA steric blockers, for example, have limited target engagement, insufficient therapeutic response, and significant off-target toxic effects. In addition, miRNA-targeting therapeutics can induce unforeseen side effects given that a single miRNA may regulate hundreds of downstream targets, resulting in complex biological responses^32,33^. Moreover, intracellular delivery of these chemically modified RNA molecules has induced adverse immunogenic responses in clinical trials, primarily through TLR and RIG receptor stimulation^4,34–36^. Lastly, while RNA knockdown only requires a low concentration of cytosolic siRNA, other applications remain limited by challenges in delivering sufficient levels of RNA due to endosomal entrapment^37^.

The discovery of diverse families of RNA-targeting CRISPR/Cas systems has opened new avenues toward refined RNA-targeting approaches that enable robust RNA degradation, editing, or steric inhibition of intracellular RNAs^38–43^. However, the broad application of RNA degradation with Cas13 nucleases is limited by collateral effects, where Cas nucleases degrade bystander RNAs upon target interaction^44–47^. Moreover, a recent study identified crRNA-independent collateral activity in major Cas13 families, raising further safety concerns for the clinical deployment of this RNA-targeting nuclease^48^.

While CRISPR discovery^40–42^ and rational engineering^49^ approaches may address these Cas13 limitations, therapeutic applications of programmable nucleases remain limited to diseases driven by overexpression of pathological RNA species. Given the diversity of disease mechanisms involving both coding and noncoding RNAs, there is a pressing need for RNA therapeutic modalities that specifically block RNA splice sites, reverse miRNA-mediated silencing, relax RNA secondary structures, and block intermolecular RNA interactions (Fig. 1a). While RNA splice donor/acceptor sites have been effectively targeted with dRfxCas13d, using this system to perturb high-affinity interactions, such as those involving RNA-protein and RNA-ribonucleoprotein (RNP) complexes remains challenging^43^.

**Fig. 1:**
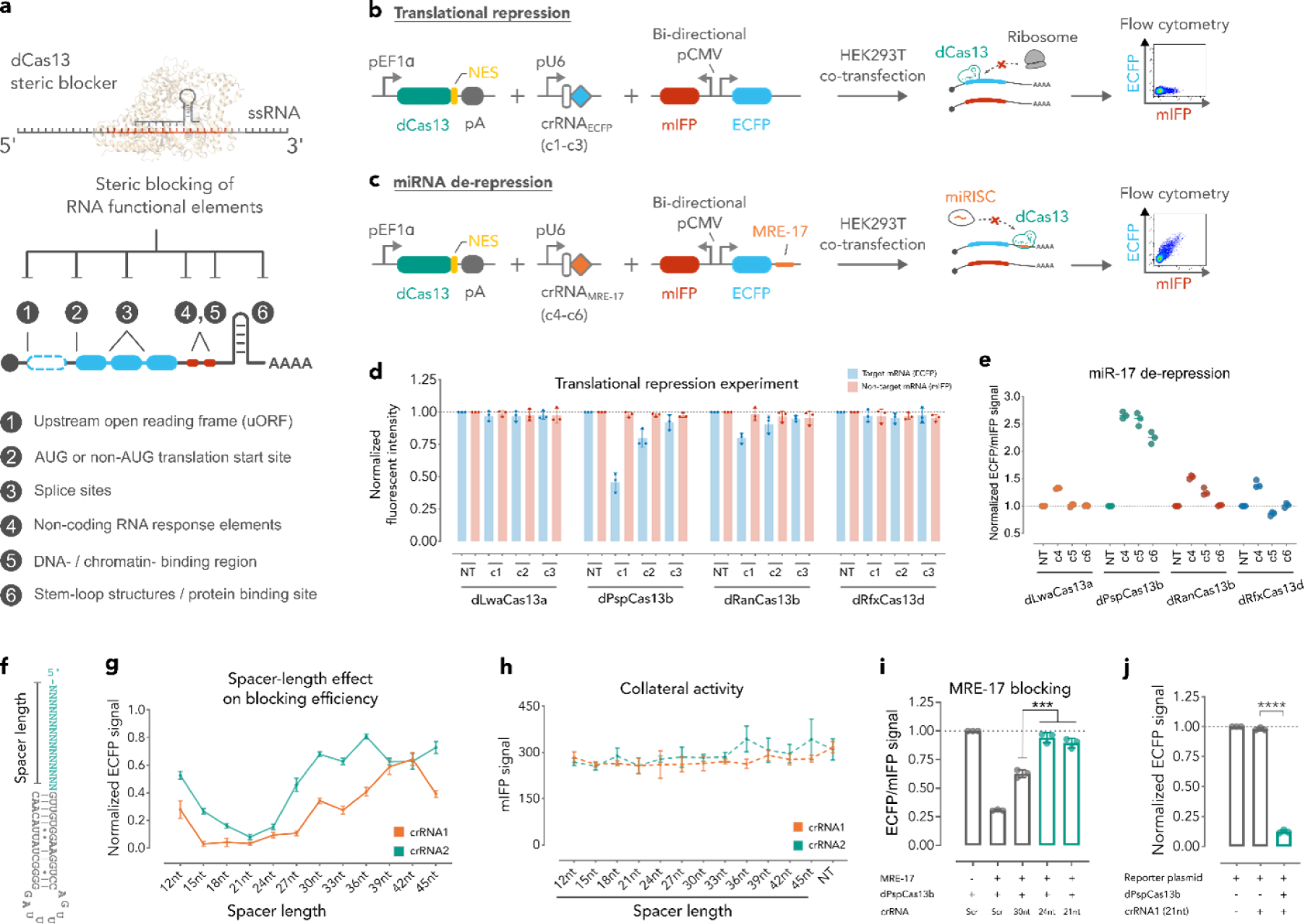
Robust and collateral activity-free RNA targeting with CRISPR-Lock 1 and CRISPR-Lock 2. **a.** Implementation of deadCas13 CRISPR RNA steric blocker (CRISPR-Lock) to sterically interfere with functional RNA elements. **b,c.** Schematic diagram of the mammalian bi-directional fluorescent reporter system used to evaluate the blocking efficiency and collateral activ-ity of dCas13 systems in the context of translational start site blocking (translational repression, **b**) or MRE blocking (miRNA de-repression, **c**). Abbreviations: NES, nuclear export signal; pA, poly-A sequence; crRNA_ECFP_, ECFP-targeting crRNAs; MRE-17, miRNA response element with full complementarity to miR-17; crRNA_MRE-17_, MRE-1-targeting crRNAs; miRISC, miRNA-induced silencing complex. **d.** Evaluation of the steric blocking efficiency of four different dCas13 orthologues, measured as relative downregulation of ECFP expression upon blocking of the ECFP translational start site. Three different crRNAs were used for each orthologue. Gating strategies are included in Supplementary Fig. 1a. **e.** Relative upregulation of ECFP/mIFP signal upon blocking of the MRE-17 located downstream of the ECFP coding sequence. **d-e.** ECFP and mIFP expression was measured by flow cytometry (n = 3) and normalised to non-targeting (NT) control group. **f.** Schematic diagram of crRNA from PspCas13b. **g.** crRNAs with shorter spacer mediate stronger blocking of the target RNA sequence, with optimal steric inhibition achieved with 18-24 nt long spacer. Blocking efficiency is measured by the relative downregulation of ECFP expression upon targeting of its translational start site. **h.** Measurement of collateral activity of dPspCas13b upon targeting highly expressed ECFP transcript, as measured by the downregulation of non-target mIFP mean fluorescent intensity signal. **i.** Short crRNAs with 24-nt or 21-nt long spacer mediate a significantly stronger blocking of MRE-17 in mammalian cells. **j.** Expression of crRNA-only in mammalian cell is insufficient to medi-ate steric inhibition, and efficient blocking requires both the expression of dPspCas13b and its short crRNA. Measurements are normalised to the no gRNA control group. In **d-e**,**g-j**, data are shown as mean *±* s.e.m., in **i,j**, *P* values are calculated by two-tailed Student’s *t*-test. ^***^*P <* 0.0001.

Here, we introduce CRISPR-Lock, a robust, high-affinity CRISPR-based RNA steric blocker. We tested four major dCas13 effector families and identified Prevotella sp. dCas13b (dPspCas13b) as the strongest candidate for effective perturbation of RNA intermolecular interactions. Subsequent optimisation of the size and design of the crRNA for dPspCas13b significantly improved its blocking efficiency. Further, we demonstrated that CRISPR-Lock can modulate protein expression after strategic design of crRNAs that target the Kozak sequence or miRNA response elements (MREs) within target RNAs. The ability of CRISPR-Lock to specifically derepress miRNA-mediated silencing through specific sites makes it an effective high-resolution perturbation tool for the systematic dissection of noncoding RNA functions. Finally, we demonstrate the therapeutic potential of CRISPR-Lock by reversing the cellular pathology of type 1 myotonic dystrophy (DM1), an RNA-dominant repeat expansion disease currently without a cure^50–52^. Using an optimised CRISPR-Lock design that facilitates delivery, we achieved robust correction of DM1-associated protein sequestration and splicing dysregulation^57^. Notably, targeting the (CUG)_n_ repeat expansion sequence with CRISPR-Lock, both in DM1 patient-derived cells and in HSA^LR^ mice *in vivo*, substantially outperformed a third-generation antisense oligonucleotide. Our results establish CRISPR-Lock as a robust and versatile RNA steric blocker with promising therapeutic prospects for treating RNA-based disorders.

## 2 Results

### 2.1 Optimisation of CRISPR-based RNA steric blocker design

To identify the optimal chassis for CRISPR-Lock, we measured the steric blocking efficiencies of different dCas13 orthologues using a bidirectional fluorescent reporter system in HEK293T cells (Figs. 1b-e). Orthologues from four major dCas13 families, each fused to a C-terminal nuclear export signal (NES), were tested alongside six crRNAs for each orthologue. crRNAs 1 to 3 (c1 – c3) targeted the 5’UTR, Kozak sequence, or the coding sequence of the ECFP transcript (Extended Data Fig. 1a). To test for translational inhibition by CRISPR-Lock, cells were co-transfected with plasmids encoding a dCas13-NES, an ECFP-targeting crRNA, and a bidirectional fluorescent reporter expressing ECFP (target transcript) and mIFP (non-target transcript) (Fig. 1b). Targeting the ECFP translational start site with dLwaCas13a-NES, dRanCas13b-NES, and dRfxCas13d-NES had minimal impact on ECFP levels. In contrast, targeting with dPspCas13b-NES moderately reduced ECFP expression (*∼*50% repression), particularly with crRNA 1, which overlaps with the Kozak sequence and immediate upstream 5’UTR sequences (Fig. 1d and Extended Data Fig. 1a). Targeting the coding sequence (CDS) of ECFP with crRNA 3 led to minimal knockdown. These results show that an efficient protein knockdown with CRISPR-Lock is achieved by inhibiting translational initiation. Furthermore, the specific downregulation of ECFP with dPspCas13b-NES did not significantly affect the non-targeted mIFP reporter, indicating that the steric-blocking mechanism of action did not cause any off-target collateral effects (Figs. 1b,d).

We next assessed the capacity of dCas13 orthologues to intervene in miRNA-mediated gene regulation by acting as miRNA target protectors (TP)^53,54^. For this purpose, we pinpointed three miRNAs - miR-17, miR-92a, and miR-222 - abundantly expressed in HEK293T cells^55^. We inserted their respective, fully complementary miRNA response elements (MREs) into the 3’UTR of an ECFP gene in our bidirectional reporter construct (Fig. 1c). As expected, this setup led to cleavage of the ECFP transcript by endogenous miRNAs, decreasing expression of the miR-17, miR-92a, and miR-222 ECFP reporters by approximately 74%, 58%, and 25%, respectively (Extended Data Fig. 1b).

To further evaluate the blocking efficiency of different dCas13 orthologues, we co-transfected HEK293T cells with plasmids encoding a bidirectional miR-17 reporter, dCas13-NES, and MRE-targeting crRNAs (crRNA 4 to 6; c4 – c6). In the case of dPspCas13b-NES, we observed 2.3-2.6-fold upregulation of ECFP expression, indicating efficient protection from miRNA degradation. Conversely, transfections with dLwaCas13a, dRanCas13b, and dRfxCas13d showed negligible changes in ECFP levels (Fig. 1e). These results imply that strong target binding is not an intrinsic property of all dCas13 proteins and may not be necessary for the native CRISPR/Cas13 functions of different orthologs. While previous studies have shown highly efficient RNA knockdown with LwaCas13a^38,56,57^, RanCas13b^58,59^, and RfxCas13d^43,60,61^, our results reveal that nuclease-deficient versions of these proteins have low target engagement, making them less suitable as candidates for potent, programmable RNA steric blockers. In contrast, dPspCas13b demonstrated superior blocking efficiency and was chosen as the chassis for our CRISPR-Lock system.

While RNA blocking with dPspCas13b-NES mediated partial silencing and mRNA protection, broader applications of this technology will require more robust steric inhibition. To improve blocking efficiency, we cloned an additional nuclear export signal (NES) at the N-terminus of dPspCas13b to improve its cytoplasmic localisation. The dual-NES design improved the protection of transcripts from miRNAs by approximately 20% (Extended Data Figs. 1c-e). Since spacer length was previously shown to affect the activity of Cas proteins in various CRISPR systems^62–64^, we sought to refine CRISPR-Lock’s steric blocking by optimising crRNA spacer length. Testing spacer lengths ranging from 12-45 nucleotides for ECFP- and MRE-targeting crRNAs, we found that shortening the spacer gradually improved blocking efficiency, with peak performance (complete repression of target gene expression) achieved using 18-24 nucleotide spacers (Figs. 1f,g and Extended Data Fig. 1f). Further truncation or extension beyond the native 30-nt length decreased the blocking efficiency (Fig. 1g). Moreover, this improved blocking did not increase off-target bystander activity (Fig. 1h and Extended Data Figs. 1f,g), underscoring CRISPR-Lock’s potential for targeted gene silencing without collateral effects. Similarly, MRE targeting with truncated crRNAs achieved 80% – 100% protection from miRNA silencing, surpassing the 50% – 60% protection achieved with crRNAs of native length (Fig. 1i and Extended Data Fig. 1g). We also observed that crRNA alone is insufficient to mediate the observed steric inhibition effect (Fig. 1j), suggesting that dPspCas13b protein significantly contributes to target-binding, likely via electrostatic and stacking interactions that complement Watson-Crick base pairing between the crRNA and its target. Our results outline a design strategy for a robust CRISPR-based RNA steric blocker capable of effectively halting trans-lational initiation and shielding RNAs from miRNA-mediated downregulation. The refined system, comprising dual-NES dPspCas13b guided by an 18-21nt spacer crRNA, is hereafter referred to as CRISPR-Lock 2.

### 2.2 High-resolution perturbation of miRNA-target interactions with CRISPR-Lock

To characterise the spatial resolution of CRISPR-Lock, we generated a set of crRNAs tiled over the target MRE (Fig. 2a). Individual assessment of these crRNAs reveals that the relative distance between the crRNA and the target MRE partially determines the efficiency of CRISPR-Lock miRNA target protector (CRISPR-Lock TP). We observed that extensive coverage by crRNAs in positions 4 – 6 (p4 – p6) was associated with higher transcript protection (Figs. 2b-d). Additionally, cover-age of the MRE at its 3’ end (p2 – p3) resulted in more efficient target protection compared to equivalent coverage at the 5’ end (p7 and p8), suggesting the importance of targeting the miRNA seed-complementary region to achieve efficient target protection (Figs. 2b-d). Full complementarity in the miRNA seed region has been considered the minimal element that enables miRNA-mediated silencing. This partially explains the improved target protection upon coverage of the 3’ end com-pared to the 5’ end of the target MRE^14,15^. Remarkably, reprogramming CRISPR-Lock to bind as close as 6-nt upstream or downstream of the MRE (p1 and p9) had no effect on ECFP expression, suggesting that perturbation of RNA interactions with CRISPR-Lock is achieved with high spatial resolution (Figs. 2b-e). We also observed that simultaneous delivery of dual crRNAs led to improved protection compared to each crRNA on its own (Fig. 2e). This strategy can be leveraged to further enhance CRISPR-Lock’s MRE-blocking efficiency.

**Fig. 2:**
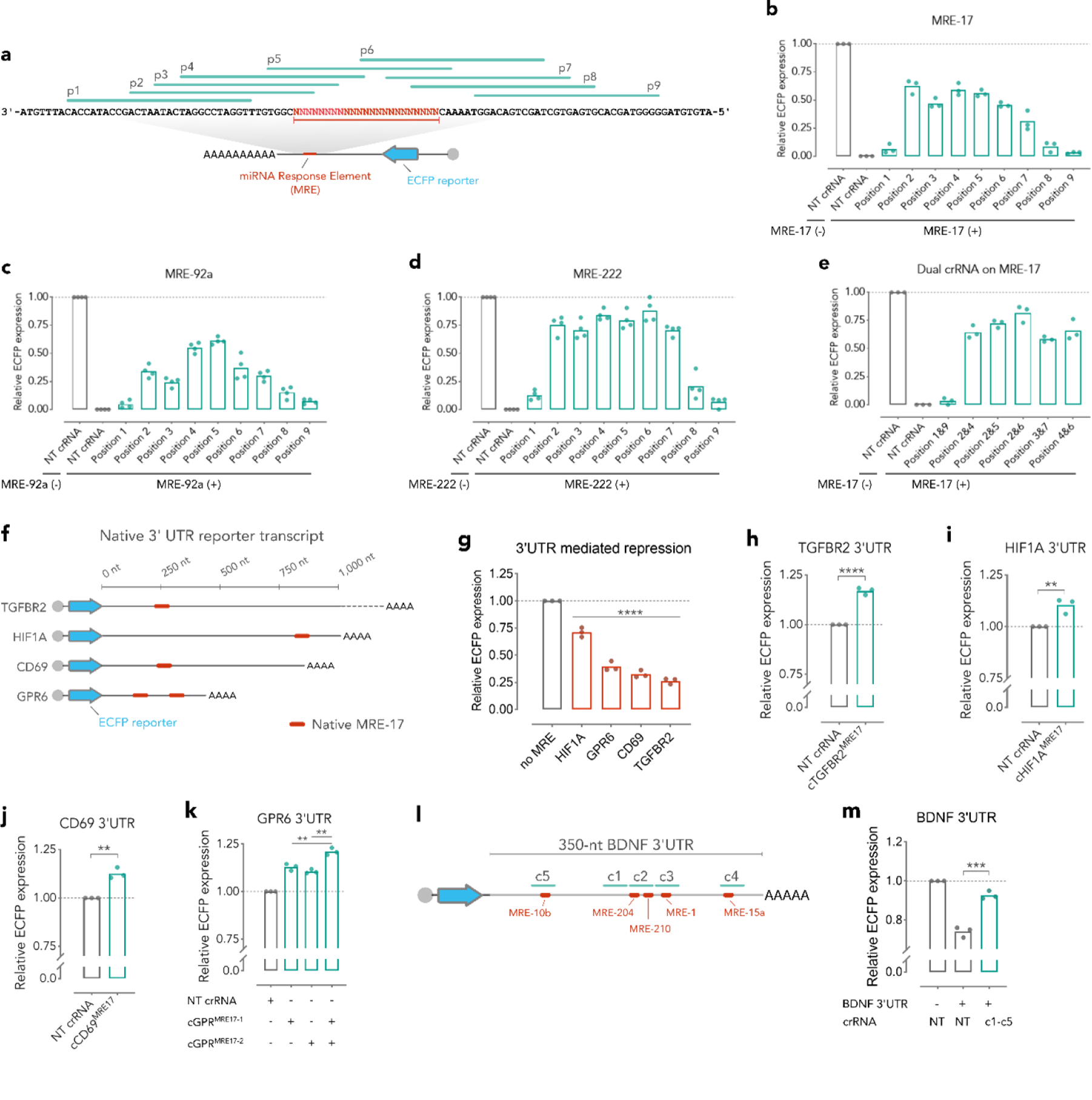
Protection of mRNA from endogenous miRNA activity and high-resolution per-turbation of miRNA response elements with CRISPR-Lock 1. **a.** Schematic diagram showing the crRNA tiling of the full complementary MRE region located downstream of the ECFP coding sequence. Nine positions of crRNA, indicated as p1 – p9, were evaluated. **b-d.** Evaluation of the spatial resolution of the steric inhibition mediated by CRISPR-Lock 1 guided by crRNAs targeting different positions surrounding the MRE as depicted in **a**. **b-d.** Reversal of miRNA-mediated silenc-ing was evaluated for three endogenous miRNAs, miR-17 (**b**), miR-92a (**c**), and miR-222 (**d**). **e.** Combination of two crRNAs located side-by-side increases the steric blocking efficiency of CRISPR-Lock. In *b–e*, measurements are normalised to the non-targeting (NT) control groups. **f.** Schematic diagram indicating the location of native MRE-17 in the 3’UTR reporter transcripts. **g.** Cloning of MRE-containing 3’UTR from various genomic loci downstream of the ECFP fluorescent reporter resulted in its downregulation. Measurements are normalised to the no MRE control group. **h-k.** Upregulation of ECFP expression after steric inhibition of native MRE-17 sequence from TGFBR2 (**h**), HIF1A (**i**), CD69 (**j**), and GPR6 (**k**) 3’UTRs with CRISPR-Lock 1, normalized to non-targeting (NT) crRNA control. **l.** Schematic diagram indicating the location of five native MREs along the fragment of BDNF 3’UTR. **m.** Multiplexed blocking of native MRE sequences in the 3’UTR of BDNF reporter resulted in significant de-repression of ECFP expression, as measured by flow cytometry and normalized to ECFP expression of the control sample. In **b-e**, **h-m** \*\**P <* 0.01; \*\*\**P <* 0.001; \*\*\*\**P <* 0.0001; In **i-j** P values are 0.0078 **(i)** and 0.0015 **(j)**. In (**g**) *P* value is calculated with one-way ANOVA with Dunnett’s multiple comparisons test on data using no MRE group as control. **h-m**, *P* values were calculated with two-tailed Student’s *t* -test.

In eukaryotic cells, near-perfect complementarity between a miRNA and its MRE leads to a robust silencing of the target mRNA via argonaute-mediated cleavage^65,66^. Meanwhile, partial miRNA: MRE complementarity results in translational inhibition followed by mRNA decay, allowing for fine-tuning of target expression levels^67^. To evaluate the efficiency of CRISPR-Lock TP in the context of mismatch-riddled MREs, we cloned four reporter plasmids containing native 3’UTRs known to harbour active MREs^68,69^. We observed 25%–75% downregulation of reporter expression in these 3’UTR reporters compared to controls, confirming the presence of functional MREs (Figs. 2f,g). Next, we co-transfected HEK293T cells with plasmids encoding a 3’UTR reporter and CRISPR-Lock components targeting the corresponding MREs. We observed that CRISPR-Lock TP mediated a 10%–17% upregulation of reporter expression in the TGFBR2, HIF1A, and CD69 3’UTR reporters (Figs. 2h-j). Moreover, we tested the GPR6 and BDNF 3’UTRs, which allowed us to target multiple MREs in a single mRNA. Targeting multiple MREs achieved significantly better protection than delivering the crRNAs individually and restored target expression levels to those achieved by deleting the whole 3’UTR (Figs. 2k-m).

Next, we sought to test the effect of the relative position between CRISPR-Lock crRNA and its target MRE. Applications of CRISPR-Lock TP to achieve precise perturbation of individual miRNA:MRE interactions are limited by the fact that any given MRE is found in hundreds of transcripts, making it challenging to design crRNAs that target a specific transcript (Extended Data Fig. 2a). We hypothesised that a particular miRNA:MRE interaction could be perturbed by using crRNAs that not only partially overlap with the target MRE, but also target immediately neighbouring sequences. This approach should discriminate between target and non-target MREs (Extended Data Fig. 2b). To assess this strategy, we designed 12 crRNAs with varying degrees of MRE coverage (Extended Data Figs. 2c,d). We observed that consecutive mismatches at the 5’ or 3’ end of the spacer impaired the efficiency of CRISPR-Lock TP, with a substantially stronger effect at the 5’ end (Extended Data Figs. 2e-h). Notably, consecutive mismatches of approximately 14nt at the 5’ end of the spacer completely abolished the CRISPR-Lock target protector activity, while the consequences of similar-sized mismatches at the 3’ end were milder (Extended Data Figs. 2e-h). In contrast, 6nt mismatches on either 5’ or 3’ end of the spacer moderately reduce CRISPR-Lock TP activity by 30%-50% (Extended Data Figs. 2e-h). These results demonstrated that the 5’ end of the crRNA is particularly important to CRISPR-Lock binding; thus, designing crRNAs that partially cover the 3’ end of the MRE and include target-specific sequences immediately downstream, substantially increases specificity by robustly blocking only the target MRE while leaving non-target MREs unperturbed. Taken together, our studies indicate a strategy for designing an effective CRISPR-Lock TP to achieve precise perturbation of individual miRNA:MRE interactions.

Next, we applied CRISPR-Lock to systematically probe RNA 3’UTRs for unbiased identification of functional MREs. We cloned segments of 3’UTR of PD-L1 (also known as *CD274*), predicted to contain multiple mRNA destabilizing elements^70^, into a reporter plasmid. This resulted in a 20%–60% downregulation of reporter expression (Extended Data Figs. 3a,b). Remarkably, a short 387nt segment of the PD-L1 3’UTR, comprising 14.3% of its total length, was responsible for approximately 75% of the silencing effect observed with the full-length 3’UTR (Extended Data Fig. 3b). To pinpoint the location of destabilising elements within the 3’UTR, we used CRISPR-Lock TP and designed 14 crRNAs to tile this segment (Extended Data Fig. 3c). Guided by crRNA-4 or crRNA-5 (c4 or c5), CRISPR-Lock achieved an 8%–15% upregulation in ECFP expression (Extended Data Fig. 3d). A subsequent tiling experiment with additional crRNAs identified a peak derepression of *∼*23% between crRNA positions c4.1 and c4.5 (Extended Data Fig. 3e). Masking this same region within the full 3’UTR reporter replicated this effect, indicating a destabilising element within the PD-L1 3’UTR covered by crRNA-4 and crRNA-5 (Extended Data Fig. 3f). Cross-referencing this region with a miRNA database revealed a sequence complementary to the miR-200b/200c/429 seed (Extended Data Fig. 3g). In this way, we demonstrated an application of CRISPR-Lock TP for unbiased identification of a functional destabilising element within the *PD-L1* 3’UTR.

### 2.3 *In vitro* correction of type 1 myotonic dystrophy (DM1) pathology

We next sought to apply CRISPR-Lock technology for the treatment of type 1 myotonic dystrophy (DM1). DM1 is an RNA-dominant disease associated with (CUG)_n_ repeat expansion in the 3’UTR of the DMPK gene. This expansion results in the toxic gain of function and the sequestration of intracellular splicing factors^71^. The resultant loss of splicing factor function causes systemic splicing dysregulation, leading to diverse symptoms, including myotonia, neurological symptoms, insulin resistance, and cardiac conduction defects^72^. Aiming to develop a robust therapeutic strategy against DM1, we developed a system to deliver CRISPR-Lock 2 components into patient-derived muscle cells to target the expanded (CUG)_n_ repeat sequence and release the sequestered splicing factors (Fig. 3a).

**Fig. 3:**
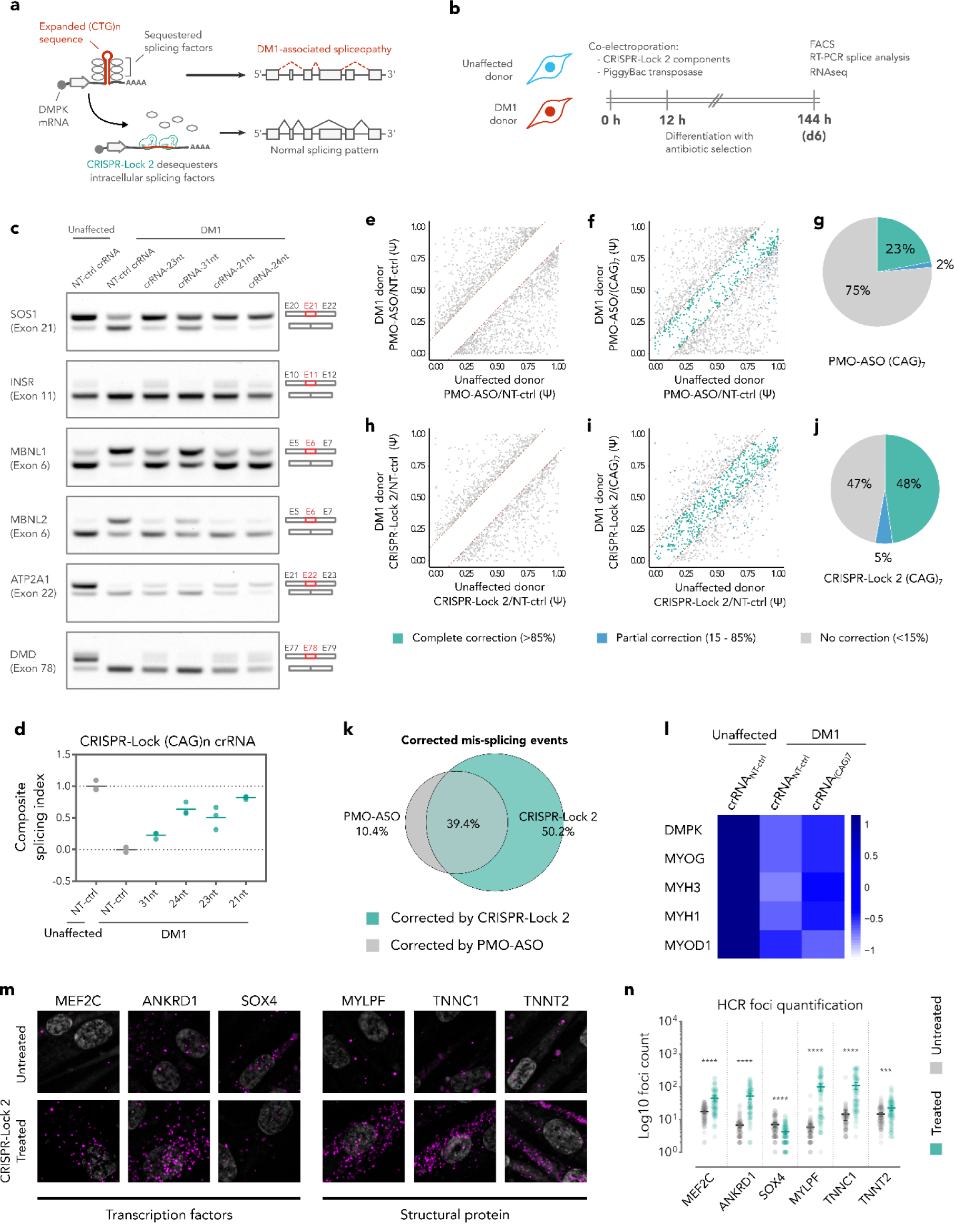
Reversal of DM1-associated pathological process in patient-derived muscle cells with CRISPR-Lock 2. **a.** Implementation of CRISPR-Lock to reverse DM1-associated pathologies by blocking repeat-expanded (CUG)_n_ sequences in the 3’UTR of DMPK mRNA, releasing splicing factors and correcting splicing dysregulation. **b.** Schematic of experimental timeline. CRISPR-Lock components were electroporated into patient-derived myoblast cells on day 0, followed by 6 days of muscle cell differentiation in tissue culture. **c.** Correction of splicing dysfunction measured by semiquantitative RT-PCR after 6 days of treatment. Uncropped gel images are included in Supple-mentary Fig. 2d **d.** Composite splicing index from a panel of 6 mis-spliced transcripts, normalized to NT and DM1 NT control groups, showing splicing correction upon CRISPR-Lock treatments. **e-f.** Scatterplot depicts splicing indices of all mis-spliced transcripts in muscle cells treated with NT-ctrl PMO-ASO (**e**) or (CAG)_7_ PMO-ASO (**f**). **g.** Pie chart showing the proportion of fully, partially, and uncorrected splicing events in unaffected DM1 cells treated with PMO-ASO. **h-i.** Scatterplot depicts splicing indices of all mis-spliced transcripts in muscle cells treated with NT-ctrl CRISPR-Lock 2 (**h**) or (CAG)_7_ CRISPR-Lock 2 (**i**). **j.** Pie chart showing the proportion of fully, partially, and uncorrected splicing events in unaffected or DM1 cells treated with CRISPR-Lock 2. **k.** Pie chart showing the pro-portion of corrected mis-spliced transcripts achieved with PMO-ASO or CRISPR-Lock 2. **l.** RNAseq evaluation of expression of *DMPK* mRNA and muscle differentiation markers in human muscle cells treated with NT-treated or (CAG)_7_ CRISPR-Lock 2. **m-n.** HCR analyses in DM1 cells expressing either (CAG)_7_ CRISPR-Lock (treated) 2 or NT CRISPR-Lock 2 (untreated). **n.** HCR quantifica-tion of genes shown in **m**. NT, non-targeting control; PMO-ASO, Phosphorodiamidate morpholino – antisense oligonucleotide; (CAG)_7_, 21-nt sequence targeting (CUG)_n_ repeat expansion. *P* values were calculated by two-tailed Student’s *t* -test; \*\*\**P <* 0.001; \*\*\*\**P <* 0.0001; n.s., not significant. Fluorescence-activated cell sorting (FACS) gating strategies are included in Supplementary Fig. 1b.

To improve the intracellular delivery of the CRISPR-Lock components, we first designed a single construct encoding both dCas13 protein and the crRNA components (Extended Data Fig.4a). This single-construct CRISPR-Lock format achieved similar blocking efficiency to the dual-construct format (Extended Data Fig. 4b-g), simplifying the delivery process. Next, we co-electroporated patient-derived unaffected (WT) and DM1 myoblasts - which contained approximately 2,600 (CTG)_n_ repeats - with a single-construct CRISPR-Lock carrying P2A-EGFP reporter and blasticidin-resistance cassettes. The plasmid also contained Super PiggyBac transposon sequences for transposase-mediated genomic integration, ensuring long-term expression of CRISPR-Lock com-ponents (Extended Data Fig. 4a). After testing four crRNA designs in WT and DM1 myoblasts (Extended Data Fig. 5a) and differentiating them into myotubes, we collected total RNA samples to measure the correction of splicing dysregulation using a panel of six DM1 severity biomarkers^73^ (SOS1 exon 21, INSR exon 11, MBNL1 exon 6, MBNL2 exon 6, ATP2A1 exon 22, and DMD exon 78) (Fig. 3b). The (CUG)_n_-targeting CRISPR-Lock 2 construct achieved approximately 85% correc-tion of the composite splicing index for these biomarkers compared to control (Figs. 3c,d, Extended Data Fig. 5b-g). These results are comparable to the correction achieved with the highest treat-ment dose of third-generation phosphorodiamidate morpholino antisense oligonucleotide (PMO-ASO) steric blockers (Extended Data Fig. 5h-k).

### 2.4 CRISPR-Lock 2 outperforms third-generation antisense oligonucleotides

To further compare CRISPR-Lock 2 and PMO-ASO treatment, we assessed the level of transcriptome-wide splicing correction achieved by these approaches. We performed RNA sequencing on total RNAs from myotubes treated with CRISPR-Lock 2, PMO-ASO, or control (Figs. 3e-j; Extended Data Figs. 6a-c). Comparing unaffected and DM1 myotubes resulted in the identification of 1,309 high-confidence dysregulated splicing events, defined by a Percent Spliced In (PSI) difference of greater than *|*0.15*|* (Figs. 3e-j, Extended Data Figs. 6a,b). The PSI values of dysregulated splicing events revealed that CRISPR-Lock 2 treatment led to notably more extensive splicing correction compared to the high-dose PMO-ASO treatment (1 *µ*M): CRISPR-Lock 2 treatment led to a 53% while PMO-ASO resulted in a 25% correction rate (Figs. 3e-j). Hierarchical clustering further showed that CRISPR-Lock 2 treatment resulted in a substantially higher level of correction compared to both low dose (250 nM) and high dose (1 *µ*M) PMO-ASO (Extended Data Figs. 6a,b). Moreover, genome browser visualisation of DM1-associated splicing factors, transcription factors, and structural proteins highlighted a more efficient correction of splicing dysregulation in CRISPR-Lock 2-treated myotubes in comparison to those treated with PMO-ASO (Extended Data Fig. 6c). Remarkably, CRISPR-Lock 2 also corrected splicing dysregulation that remained unaltered by PMO-ASO, without degradation of the DMPK transcripts, supporting the notion that CRISPR-Lock 2 operates via a steric blocking mechanism (Figs. 3k,l, Extended Data Figs. 7a,b). Finally, differential expression analysis indicated that correction of splicing dysregulation in the patient-derived myotubes was followed by an improvement in DM1-associated differentiation defects (Figs. 3l-n, Extended Data Figs. 7c-f). For instance, CRISPR-Lock 2-treated myotubes displayed an upregulation of late muscle differentiation markers such as MYOG and MYH and a downregulation of early-differentiation markers (MYOD), which was not observed in control or PMO-ASO treated cells (Fig. 3l, Extended Data Fig. 7f). Hybridization chain reaction (HCR) experiments validated the correction in the expression levels of key muscle transcription factors (*MEF2C*, *ANKRD1*, and *SOX4*) and structural proteins (*MYL11*, *TNNC1*, *TNNT2*) in cells treated with CRISPR-Lock 2 (Figs. 3m-n, Extended Data Figs. 7c-e). In summary, these data indicate the therapeutic potential of CRISPR-Lock 2 to correct the complex molecular pathology of DM1.

### 2.5 Design of compact single-AAV CRISPR-Lock 2 and *in vivo* correction of DM1 pathology

Clinical translation of CRISPR-Lock 2 requires safe and efficient delivery methods for targeting therapeutically relevant tissue types. Adeno-associated viruses (AAVs) have been widely used to deliver therapeutic cargo, such as approved drugs, and are a popular choice in clinical trials due to their tissue customizability and favourable safety profile. At *∼*5.1 kilobases (kb), the CRISPR-Lock 2 construct does not fall within the *∼*4.7 kb cargo size limit of AAV (Fig. 4a). Therefore, we first attempted to generate a smaller therapeutic construct by using a shorter EFS promoter. However, the expression levels achieved with the EFS promoter were not sufficient for mediating the correction of splicing dysregulation *in vivo* (Extended Data Figs. 8a-c). An alternative solution, the dual-AAV approach, has widely been used to deliver large CRISPR cargos. However, this approach can require higher treatment doses, increasing the potential adverse effects related to the treatment^74^.

**Fig. 4:**
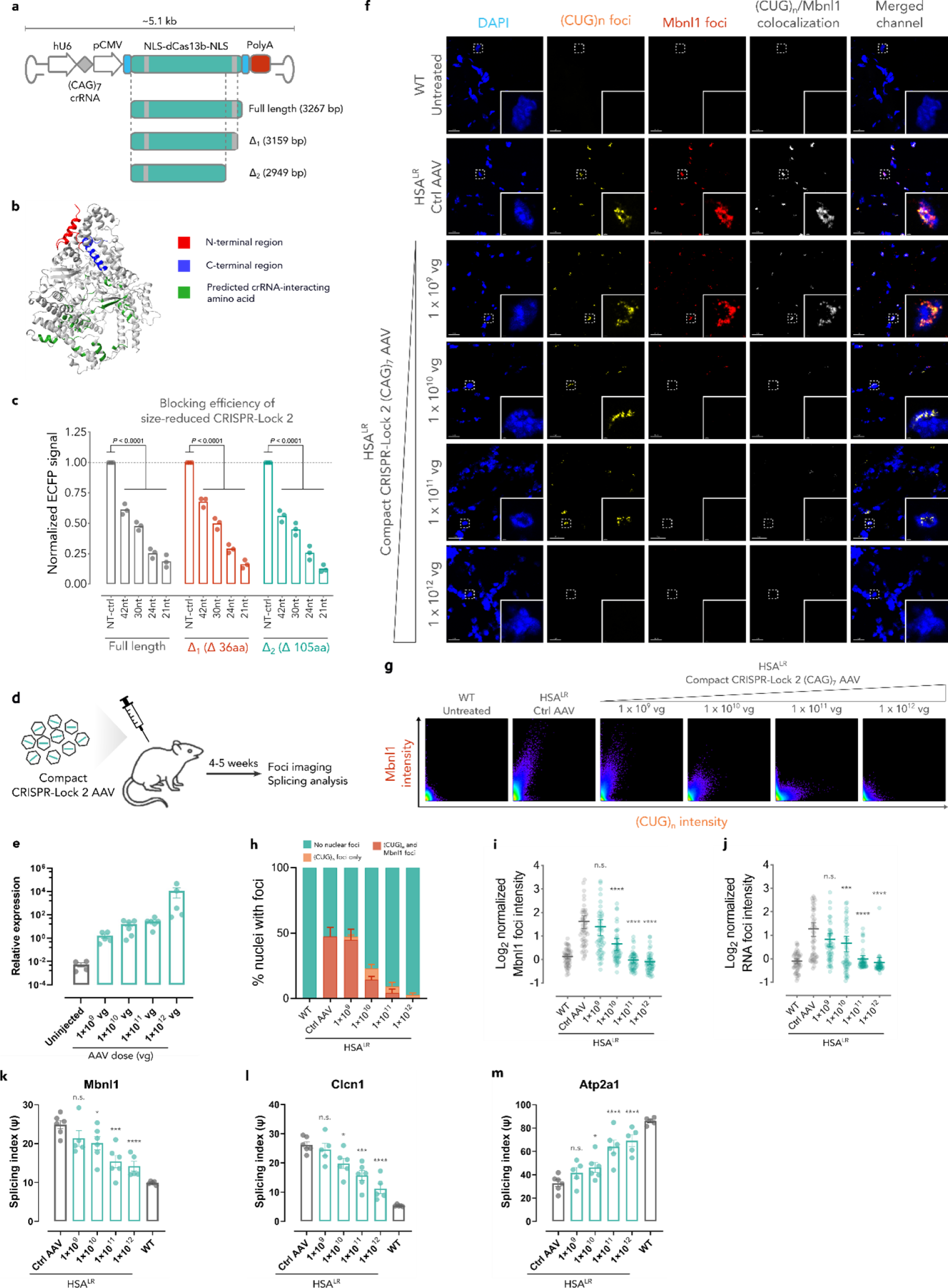
Compact CRISPR-Lock 2 enables all-in-one AAV design and dose-dependent correction of DM1-associated pathologies in HSA^LR^ mice. **a-c.** Reducing the size of the CRISPR-Lock 2 system enables the packaging of all its components into a single AAV construct while maintaining its steric-blocking efficiency. **a.** Design strategy to enable single-AAV packaging of CRISPR-Lock 2 components. **b.** AlphaFold 2 prediction of dPspCas13b structure in ribbon representation. The N-terminal region (red), C-terminal region (blue), and predicted crRNA-interacting amino acids (green) are highlighted. **c.** Shortened dPspCas13 proteins have similar steric blocking efficiency compared to the full-length protein. This assay was performed on HEK293T cells using a bidirectional reporter system, and ECFP signal was measured with flow cytometry, measurements normalised to non-targeting (NT) control group. *P* values were calculated with one-way ANOVA with Dunnett’s multiple comparisons test on data using NT control groups as control. **d.** Intramuscular (IM) injection of WT or HSA^LR^ mice with single-AAV CRISPR-Lock 2 construct. Tibialis anterior (TA) muscles of these mice were injected with varying dose of AAV-packaged CRISPR-Lock 2 guided with either NT control crRNA or (CAG)_7_ crRNA. **e.** Expression level of CRISPR-Lock 2 system in the TA muscles of injected HSA^LR^ mice, normalised to GusB, as measured with ddPCR. **f.** Dose dependent elimination of DM1-associated Mbnl1 and (CUG)_n_ nuclear foci in HSA^LR^ muscle tissue after treatment with single-AAV CRISPR-Lock 2. Repeat-expanded (CUG)_n_ foci are labelled with FISH (Cy3 channel), Mbnl1 foci were labelled with immunofluorescent (AF647 channel), nuclei are labelled with DAPI. **g.** Scatterplot showing the voxel intensity of the Cy3 and AF647 channel, representing (CUG)_n_ and Mbnl1 respectively. In **f-g**, images are representative of n *≥* 3 independent replicates. **h.** Dose dependent reduction of the number of nuclei containing either both DM1-associated (CUG)_n_ and Mbnl1 foci or only (CUG)_n_ foci in TA muscle tissues of mice treated with single-AAV CRISPR-Lock 2 or controls. **i-j**, Dose dependent reduction of mean intensity of DM1-associated Mbnl1 foci (**i**) and (CUG)_n_ foci (**j**) in TA muscle tissues of mice after treatment with single-AAV CRISPR-Lock 2 or controls. In **i-j**, n = 50 nuclei were analysed, and and P values were calculated with one-way ANOVA with Dunnett’s multiple comparisons test using Ctrl AAV groups as control values were calculated with one-way ANOVA with Dunnett’s multiple comparisons test using Ctrl AAV groups as control. \*\*\**P <* 0.001; \*\*\*\**P <* 0.0001; n.s., not significant. **k-m**, Dose-dependent correction of DM1-associated mis-splicing post single-AAV CRISPR-Lock 2 treatment in *Mbnl1* (**k**), *Clcn1* (**l**), and *Atp2a1* (**m**), as measured with ddPCR. Data are mean *±* s.e.m.

Therefore, we opted to develop a single AAV construct encoding a compact CRISPR-Lock 2 driven by a strong mammalian promoter. Protein alignment and structural prediction of dPspCas13b using AlphaFold 2 showed that the HEPN2 domain was located far away from the side channel where crRNA-target interactions occur, suggesting that this region is not directly involved in crRNA/target binding (Fig. 4b, Extended Data Figs. 8d-e), in line with previous observations^58,75^. We therefore hypothesised that while the truncation of the HEPN2 domain of dPspCas13b would significantly reduce the size of the CRISPR-Lock 2 construct, it would not sacrifice its steric blocking efficiency. We tested two truncated variants of dPspCas13b, Δ1 and Δ2, which reduce the size of the full-length dPspCas13b by 4% and 10%, respectively (Fig. 4a). The steric blocking efficiencies of these truncated variants were comparable to the full-length CRISPR-Lock system and responded similarly to short crRNAs (Fig. 4c). Therefore, the Δ2 variant was used for subsequent *in vivo* experiments and referred to as compact CRISPR-Lock 2.

Next, we sought to assess the efficacy of the compact CRISPR-Lock 2 RNA steric blocker *in vivo* using a mouse model of DM1. We used HSA^LR^ mice, an established model of DM1 which expresses a human skeletal actin gene containing a (CTG)n repeat of approximately 660–750 bp in its 3’UTR^76^. These mice exhibit molecular and symptomatic hallmarks of DM1, including protein sequestration, widespread splicing dysregulation, muscle weakness, and muscle degeneration^76,77^. Splicing dysfunction in DM1 primarily affects skeletal muscle, making this tissue our primary target for the delivery of CRISPR-Lock 2. We generated compact CRISPR-Lock 2 AAV encoding pU6-driven crRNA(CAG)_7_ and pCMV-driven compact dPspCas13b and performed intramuscular (IM) injections of 1×10E9 to 1×10E12 vector genomes (vg) of AAV into the tibialis anterior (TA) muscle of HSA^LR^ mice. We collected muscle tissues for analysis 4 to 5 weeks after injection (Fig. 4d). The compact design of CRISPR-Lock 2 allowed us to use a strong promoter, pCMV, achieving approximately 800-fold higher expression levels of the system compared to pEFS-driven full-length CRISPR-Lock 2 (Fig. 4e, Extended Data Fig. 8b).

In mice treated with compact CRISPR-Lock 2 AAV, we observed a dose-dependent release of *Mbnl1* from (CUG)_n_ RNA foci, with effects at doses as low as 1×10E10 vg (Figs. 4f,g; Extended Data Fig. 8f). Moreover, administering a higher AAV dose resulted in a reduction in (CUG)_n_ RNA foci count and intensity (Fig. 4f,g; Extended Data Fig. 8g), suggesting that desequestration of proteins from the mutant DMPK RNAs contributes to transcript destabilisation, consistent with previous studies^78^. Concurrently, we observed a dose-dependent decrease in the proportion of nuclei containing RNA-protein foci (Fig. 4h) and a significant reduction in the average intensity of both Mbnl1 and (CUG)_n_ foci within nuclei (Figs. 4i,j) in treated mice compared to controls. Notably, the reduction in foci number and intensity correlated with a dose-dependent correction of splicing dysregulation in Mbnl1 exon 5, Clcn1 exon 7, and Atp2a1 exon 22 transcripts (Figs. 4k-m), confirming the recovery of free Mbnl1 following treatment with (CUG)_n_-targeting compact CRISPR-Lock 2 AAV. Taken together, these findings support the steric blocker mechanism of action of the CRISPR-Lock system, whereby competitive binding to the expanded (CUG)_n_ repeat sequence reverses Mbnl1 sequestration, correcting the broader splicing errors that underlie the symptoms associated with DM1 in these tissues.

## 3 Discussion

The potential of RNAs as therapeutic targets was demonstrated in pioneering work by Zamecnik and Stephenson, who achieved inhibition of Rous Sarcoma Virus (RSV) replication by targeting viral 35S RNA with synthetic nucleotides^79^. Since then, RNAs have been implicated in a wide range of pathological processes, along with the development of modalities to degrade, block, edit, and replace RNAs in diseased cells^21–31,39,43,58,80,81^. Despite the regulatory approval of various RNA-targeting therapeutics^1,3,22^, a robust, genetically encoded modality capable of manipulating a wide array of coding and noncoding RNA functions remained beyond reach. Here, we have developed an RNA-targeting CRISPR-Lock system and optimized it to target protein translation, disrupt an aberrant RNA-protein interaction, and interfere with non-coding RNA (ncRNA) function.

Specifically, we identified dPspCas13b as the most effective chassis for an efficient RNA-steric blocker, which we named CRISPR-Lock. During our screening experiments, we noted that most of the tested CRISPR/dCas13 orthologues possessed relatively low blocking efficiency, indicating that high target binding affinity is not common among CRISPR/Cas13 family members despite their robust nuclease activities^38,43,82^. Previous studies have suggested that target binding and nuclease activation are decoupled in Cas13 family members, implying that robust target degradation does not require high binding affinity^83^. This apparent contradiction likely evolved since Cas13 ordinarily induces cellular dormancy in phage-infected prokaryotes. Here, Cas13 nucleases use crRNA guides to locate viral transcripts and trigger indiscriminate cleavage of both target and bystander RNAs, resulting in cellular dormancy and abortion of the viral infectious cycle^84,85^. In this context, low target binding affinity serves as a ‘safety switch’, allowing Cas13 to dissociate from the viral transcript, thus facilitating its deactivation.

Previous studies have shown that Cas13 collateral activity targets bystander RNAs through separate catalytic sites^49,56,60,75,86^. However, our experiments show that H133A/H1058A mutations in PspCas13b, which inactivate the ‘on-target’ catalytic sites, are sufficient to abolish both ‘on-target’ and collateral RNA cleavage activities, thus establishing dPspCas13b as a collateral-free RNA steric blocker.

To generate a more robust CRISPR-Lock, we explored the design space of dPspCas13b crRNA to maximise its steric blocking efficiency. Consistent with previous findings in DNA- and RNA-targeting CRISPR systems, altering the length of the spacer region in CRISPR sgRNAs significantly affects the binding affinities of Cas nucleases, which often directly impacts their nuclease activities^62–64^. Interestingly, we observed that supplying dPspCas13b with shorter crRNAs substantially improves blocking efficiency, implying that a shorter spacer increases the binding affinity of the system. A previous study revealed that conformational changes in the HEPN1 and HEPN2 domains of Cas13b are necessary to enable the entry of the target RNA into the side channel of the protein^75^. This study also found that the crRNA spacer region occupies the Cas13b side channel before target binding and partially obstructs HEPN1/HEPN2 conformational changes. Together with these observations, data collected in our experiments suggest that shortening the crRNA spacer may facilitate blocking efficiency by enabling more sterically favourable binding. Our optimized design, CRISPR-Lock 2, thus utilized a short crRNA and dual localisation signals. We demonstrated that CRISPR-Lock 2 enabled the complete silencing of strong transgene expression and protected target transcripts from degradation by the most active miRNA species in HEK293T cells.

Derepression of target RNAs from miRNA-mediated silencing can be accomplished by masking the miRNA response element, a method known as miRNA target protector^53,54^. Here, we demonstrated that CRISPR-Lock can be used as a genetically encoded miRNA target protector. Our crRNA tiling experiments outlined the design rules for crRNA positioning relative to the target MRE. The efficiency of mRNA protection with CRISPR-Lock TP correlates with the degree of MRE coverage, especially when including the seed complementary region, in line with previous observations highlighting the importance of miRNA seed region^87–90^. Coverage of immediate MRE-neighbouring sequences by directing CRISPR-Lock to bind approximately 6 nucleotides upstream or downstream of the MRE does not lead to notable mRNA protection, suggesting that CRISPR-Lock mediates protection from miRNAs with high spatial resolution. Considering that the rate of protein translation from mRNAs is often regulated by a cluster containing multiple MRE species^91,92^, high-resolution perturbation with CRISPR-Lock TP allows for the precise investigation of individual MRE functions within a regulatory cluster.

In cells, MREs from a single miRNA species can be present in hundreds of mRNAs^14,93,94^, posing a challenge for the implementation of a highly specific miRNA target protector. By characterising the mismatch tolerance of the system, we found that mismatches at the 5’ end of the crRNA spacer region substantially reduce CRISPR-Lock blocking efficiency. Therefore, designing crRNA to cover the seed complementary sequence and its adjacent upstream region would improve the targeting specificity of the CRISPR-Lock TP while maintaining robust perturbation.

miRNAs are a well-characterised class of noncoding RNAs that play crucial roles in physiological and pathological processes^5,95–104^. Aberrant miRNA expression or loss of cognate MREs has been linked to a range of malignant diseases^14,70,93,105^, making miRNAs appealing therapeutic targets. Although several miRNA-targeting therapeutic modalities exist, most aim to achieve miRNA knockouts, which can affect hundreds of downstream targets and may lead to adverse effects^102,106–109^. Our work with CRISPR-Lock demonstrates the precise perturbation of individual miRNA-target interactions by shielding the MRE, enabling the design of gene therapies that block pathological miRNA-target interactions while preserving the physiological roles of the miRNA.

Demonstrating the therapeutic potential of CRISPR-Lock, we show that this technology can be used to block pathological RNA-protein interactions in type 1 myotonic dystrophy. In contrast, previous efforts to develop a dCas13-based RNA blocker for the treatment of DM1 did not result in the correction of splicing dysregulation, partially due to inadequate blocking efficiency^47^. Congenital DM1 (CDM), the most severe clinical subtype caused by extremely long repeat expansions (*>*1,000 repeats), is characterised by distinct clinical features, including severe muscle weakness, which often compromises respiratory functions and results in high mortality rate^110–112^. Extensive investigations into DM1 therapeutics have brought candidate medicines into pre-clinical studies and clinical trials, including small molecules^113,114^, ASOs^115,116^, siRNAs^117^, RNA-targeting PIN-dCas9^118,119^, and decoy RNA-binding protein^120^. While these technologies have the potential to profoundly alter the trajectory of newborns with CDM, each therapy has limitations, including a narrow therapeutic window, open questions regarding toxicity, DSB-related chromosomal rearrangements, and collateral off-target effects, highlighting the importance of continued development of new genetic therapies^44,45,121,122^.

Our study describes the optimisation of CRISPR-Lock crRNA design for targeting DM1-associated repeat expansion. We used patient-derived myoblast expressing *∼*2,600 CUG repeat as a disease model known to appropriately recapitulate molecular and phenotypic hallmarks of the disease^123^. We show that treatment with an optimised CRISPR-Lock 2 design achieves transcriptome-wide correction of splicing dysregulation, with approximately 85% composite correction of clinically relevant splicing biomarkers. Remarkably, the reduction in splicing dysregulation achieved by CRISPR-Lock 2 outperforms that achieved with PMO-ASO, a third-generation antisense oligonucleotide with improved binding affinity and stability used in the clinic^124–126^. Beyond reducing splicing dysregulation, we observed that treatment of DM1 tissues with CRISPR-Lock 2 upregulated late myotube differentiation markers (MYOG, MYH, MEF2C, TNNT2) and downregulated early differentiation markers (MYOD, SOX4). These results suggest the activation of the late-stage muscle differentiation program and reversal of muscle maturation arrest - the phenotypic hallmark of DM1^127–129^.

Upon demonstrating the *in vitro* therapeutic potential of CRISPR-Lock 2, we sought to apply this technology to reverse DM1-associated splicing dysregulation in HSA^LR^ mice, an established preclinical model of DM1 expressing 250 CUG repeats driven by hACTA1 promoter^76^. To this end, we developed a compact CRISPR-Lock 2 by removing the predicted HEPN2 domain of dPspCas13b, enabling the packaging of all CRISPR-Lock 2 components in a single AAV construct. Importantly, this miniaturisation of the CRISPR-Lock 2 system was achieved without sacrificing the blocking efficiency of the system. Using an AAV-encoded compact CRISPR-Lock 2, we performed a dose escalation study in HSA^LR^ mice, which is an important part of evaluating the clinical potential of gene therapy. A recent trial demonstrated the safety of intramuscular delivery of 5x10E10 to 2.5x10E12 vg/kg AAV constructs in human participants. Our studies in mice involved dose ranges between 5x10E10 to 5x10E13 vg/kg. We observed MBNL desequestration, RNA foci destabilization, and correction of splicing dysregulation at doses as low as 5x10E11 vg/kg.

Our study outlines the design and application of CRISPR-Lock systems *in vitro* and *in vivo*. We demonstrated that CRISPR-Lock is robust miRNA target protector, highlighting its advantages over conventional miRNA knockdown and morpholino TPs. The study successfully demonstrated the efficacy of CRISPR-Lock in blocking pathological RNA-protein interactions in both patient-derived cells and a mouse model of type 1 myotonic dystrophy, achieving superior outcomes compared to third-generation antisense oligonucleotides (ASOs). The wider therapeutic application of this technology hinges on investigating alternative delivery methods, such as AAV vectors with enhanced muscle transduction^130,131^, eVLPs^132^ or nanoparticles, alongside optimising injection frequencies and durations to maximise splicing correction. Finally, combining CRISPR-Lock treatment with small molecules or ASOs could significantly improve therapeutic correction and facilitate phenotypic recovery *in vivo*.

## 4 Methods

### 4.1 Plasmids, primers, and construct assembly

Plasmids used in this study are described in Supplementary table 1; new plasmids generated during this study will be deposited in Addgene. All crRNA and gRNA sequences used in this study are described in Supplementary table 2. For plasmid cloning, vector backbone digestion was performed using restriction enzymes in the manufacturer’s recommended buffers. Subsequently, digested vectors were treated with 5 units of Antarctic Phosphatase (New England Biolabs) for 30 min at 37°C. Post-digestion purification of all vectors was performed using the Qiagen gel extraction kit (Qiagen) or a Qiagen MinElute PCR purification kit (Qiagen), as appropriate. When gel extraction was required, purifications were performed as per the manufacturer’s instructions but with Qiagen MinElute PCR purification columns instead of Qiaquick columns and an elution volume of 10 *µ*L using buffer EB. Vector backbones were purified with the MinElute PCR purification kit according to the manufacturer’s instruction. Inserts were generated either by PCR (for large inserts) or by annealing oligonucleotide duplexes (for crRNA cloning). PCR-amplified inserts were purified prior to their ligation into vectors using either a standard Qiagen gel extraction method or a Qiaquick PCR purification protocol, as appropriate. Ligation reactions used either 200 or 400 units of T4 DNA ligase (New England Biolabs) in the T4 DNA ligase buffer, maintaining the vector to insert molar ratio between 1:1 and 1:10. These reactions were incubated at room temperature for 10 min or at 16°C overnight, followed by a 2 min cooling on ice. Subsequently, 1–3 *µ*L of the ligation mixture was mixed with 10–50 *µ*L of Subcloning Efficiency™ DH5*α*™ Competent Cells (ThermoFisher Scientific), maintaining a volume ratio of ligation product to bacteria below 1:10. Cells were then incubated on ice for 30 min, heat shocked at 42°C for 30 s in a water bath, then 200–500 *µ*L S.O.C. medium (home-made) was added. This was followed by a 1 h incubation at 37°C with shaking. The resultant cultures were plated on LB agar plates containing ampicillin (100 *µ*g/mL), chloramphenicol (25 *µ*g/mL), or kanamycin (50 *µ*g/mL), as appropriate. Plates were incubated at 37°C overnight. Individual colonies were then picked into 3 – 8 mL antibiotic-supplemented LB media and incubated again overnight at 37°C with shaking. Plasmid purification was performed using Qiaprep Spin Miniprep columns (Qiagen) according to the manufacturer’s protocols, and the integrity of the plasmids was confirmed through diagnostic digests and sequencing with the appropriate primers.

Vectors encoding dCas13 were generated by inserting PCR amplicons containing a dCas13 orthologue, protein localization signals, and/or fluorescent markers, into a backbone vector under the control of either an EF1-alpha or CMV promoter. PCR amplifications were performed with Phusion^®^ High-Fidelity PCR Master Mix with GC Buffer (New England Biolabs) with 500 nM primers. The amplification protocols were as follows: initial denaturation at 98°C for 60 s, 40 cycles of denaturation at 98°C for 10 s, annealing under optimal conditions determined by the NEB Tm calculator for 30 s, extension at 72°C for 60-180 s, and a final extension at 72°C for 5 min. crRNA-encoding vectors were constructed by annealing oligonucleotide duplexes (IDT), followed by ligation of the resultant double-stranded fragments into vectors containing a hU6 promoter and a pol III termination signal. For all oligo annealing reactions, a mixture of 10 *µ*L of each of the forward and reverse oligos (10 *µ*M), 1× T4 DNA ligase buffer, and five units of T4 Polynucleotide Kinase (NEB) were incubated at 37°C for 30 min, ramped to 95°C for 5 min then cooled to 25°C at a rate of 0.1°C/s. The annealing reaction products were subsequently used as inserts in cloning reactions as appropriate. All new plasmids generated in this study will be made publicly available on Addgene upon publication of this study.

### 4.2 Cell culture and transfection of HEK293T cells

HEK293T cells, obtained from the American Type Culture Collection (ATCC-CRL-11268), were cultured in Dulbecco’s Modified Eagle Medium (DMEM, Gibco) containing 10% FBS (E.U.-approved, South America origin, Gibco). Cells were maintained in a 37°C incubator with 5% CO_2_ and sub-cultured upon reaching 90%-95% confluency with dilution ratios between 1:3 to 1:10. All cells tested negative for mycoplasma at least every 6 months using a VenorGeM^®^ Mycoplasma Kit (Minerva Biolabs) according to the manufacturer’s instructions.

For transfection experiments, mycoplasma-free HEK293T cells were revived from liquid nitrogen cryopreservation and were passaged at least three times prior to any experiments. For each transfection experiment, 125,000 cells were plated per well in 24-well plates (Corning) in 500 *µ*L of DMEM containing 10% FBS. Transfections were performed the next day when cells reached 70%-90% confluence. Immediately before transfection, culture medium was replaced with 500 *µ*L of DMEM containing 2% FBS and cells were incubated at 37°C and 5% CO_2_ for 30 min. Concurrently, a transfection mixture was prepared by combining plasmid DNA, Opti-MEM™ (Gibco), and polyethylenimine (PEI, Sigma). Except stated otherwise, for each well, 500 ng of dCas13-encoding plasmid, 1 *µ*g of crRNA-encoding plasmid, and 50 ng of a bidirectional reporter plasmid were diluted to a final volume of 50 *µ*L in Opti-MEM™ (Gibco) containing 1.5 *µ*g PEI. This mixture was thoroughly vortexed for 10 s, allowed to rest at room temperature for 20 min, and then applied to the cells in a dropwise manner. Following a 24 h incubation period at 37°C with 5% CO_2_, the transfection medium was replaced with fresh DMEM containing 10% FBS. After an additional 24 h (totalling 48 h post-transfection), the cells were harvested for flow cytometry analysis.

### 4.3 Flow cytometry and assessment of CRISPR-Lock blocking efficiency in HEK293T cells

Adherent HEK293T cells were harvested using 0.05% Trypsin-EDTA solution (ThermoFisher Scientific). Post-trypsinization, the cells were washed in PBS twice, and then resuspended in their respective growth medium. Before flow cytometry analysis, cells were filtered through a 70 *µ*m cell strainer (Corning) to ensure a uniform single-cell suspension. The flow cytometry measurements were conducted using the BD LSRFortessa™ cell analyser at the Weatherall Institute of Molecular Medicine Flow Cytometry Core Facility. ECFP was measured following 405 nm excitation using a 450/50 bandpass filter. EGFP was measured using 488 nm excitation with a 530/30 bandpass filter. mIFP was measured using 640 nm excitation with a 670/14 bandpass filter. Flow cytometry data were analysed with FlowJo software (Tree Star).

### 4.4 Human myoblast cell culture, electroporation, differentiation, and FACS sorting

Myoblast cell lines from healthy donors were acquired from the MRC Centre for Neuromuscular Diseases (CNMD) Biobank London (Catalog No. L954/1284 M-I) and DM1 myoblast cell line with 2,600 CTG repeats were generously provided by Denis Furling^123^. Maintenance of all myoblast cultures was carried out in Skeletal Muscle Cell Growth Medium (SkMC, PromoCell) with the addition of SupplementMix (PromoCell) and incubated at 37°C and 5% CO_2_. Cells were sub-cultured upon reaching a confluency of 90%-95%, with dilution ratios ranging from 1:3 to 1:10. All cell lines were tested for mycoplasma contamination at least every 6 months using the VenorGeM^®^ Mycoplasma Detection Kit (Minerva Biolabs), according to manufacturer’s instructions. For myoblast electroporation experiments, myoblast cells were detached using 0.05% Trypsin-EDTA (ThermoFisher Scientific) and subjected to electroporation using the Neon Transfection System kit (ThermoFisher Scientific), following the guidelines provided by the manufacturer. For the procedure, each batch of 150,000 cells was transfected with 100 ng of the Super PiggyBac Transposase Expression Vector (System Biosciences) and 600 ng of vectors encoding dCas13 and crRNA. The electroporation parameters were set at 1750 V, 10 ms, and 3 pulses. Following electroporation, cells were cultured in SkMC Growth Medium with SupplementMix (PromoCell) at 37°C and 5% CO2 for 8 – 12 h to facilitate cell adhesion to the culture plate surface. After this period, the growth medium was replaced with SkMC Differentiation Medium (PromoCell), also supplemented with SupplementMix (PromoCell), and the cells were incubated overnight. The following day, the differentiation medium was refreshed, this time incorporating 300 *µ*g/mL blasticidin (Life Technologies) to the mixture. This antibiotic-supplemented differentiation medium was renewed every 1 to 2 days over a span of 6 days to promote the differentiation of patient-derived myoblasts into myotubes.

For fluorescent-activated cell sorting (FACS) of CRISPR-Lock expressing myotubes, cells were detached using 0.05% Trypsin-EDTA (ThermoFisher Scientific). Following TE treatment, the cells were washed twice with PBS, then resuspended in SkMC media containing SupplementMix (PromoCell) and 0.1 *µ*g/mL DAPI for nuclear staining. The prepared cell suspensions were filtered through a 70 *µ*m cell strainer (Corning) to ensure a uniform single-cell suspension. Cell sorting was conducted using the BD FACSAria™ Fusion cell sorter at the Weatherall Institute of Molecular Medicine Flow Cytometry Core facility. DAPI-stained nuclei were detected by excitation at 405 nm and emission collected through a 450/50 nm bandpass filter, while EGFP was measured using 488 nm excitation with a 530/30 bandpass filter. Following sorting, cells were collected in SkMC growth medium (PromoCell) and were immediately processed for RNA extraction.

### 4.5 Extraction of human myoblast total RNA and semiquantitative PCR analyses

Total RNAs were extracted from FACS-sorted patient-derived myotubes using either the RNeasy Mini Kit (Qiagen) or the PureLink RNA Mini Kit (Life Technologies), according to the manufacturer’s instruction. The isolated total RNA samples were then subjected to DNase treatment using the TURBO DNA-free™ Kit (ThermoFisher Scientific) in a total volume of 30 *µ*L, as per the manufacturer’s protocol, to eliminate any remaining plasmid DNA. For each RNA sample, complementary DNA (cDNA) was synthesized from 75 ng of total RNA utilizing the SuperScript™ First-Strand Synthesis System (Life Technologies). A volume of 1 *µ*L of the resultant cDNA was then used in semi-quantitative PCR analyses, using DreamTaq Green PCR Master Mix (ThermoFisher Scientific) with 500 nM primers. Between 1-10 *µ*L of PCR product were then run on a 2% agarose gel and nucleic acids visualized with GelRed^®^ (Biotium). Gel documentation was performed with a BioRad GelDoc™ XR+ imager, setting exposure times to just under the threshold for pixel saturation (approximately 0.75 seconds). Subsequently, image processing was done with ImageJ, which included background subtraction and the quantification of density for both upper and lower bands of each sample. The percentage splicing inclusion (PSI, Ψ) for exon-skipping events was calculated by determining the ratio of the upper band’s density to the total band density for each gene of interest. The composite splicing index (composite PSI) was calculated as the average of corrections of each assessed transcript for each sample, considering the mean PSI values from unaffected donor cells as maximum and the mean PSI values from NT-crRNA treated sample as minimum.

### 4.6 RNA-seq library preparation and sequencing

Following total RNA extraction and DNase treatment, the RNA integrity of all specimens was assessed using either the Bioanalyzer 2100 system (Agilent Technologies) or the TapeStation 2200 system (Agilent Technologies). Only RNA samples with an RNA Integrity Number (RIN) above 8.0 were selected for library construction using the KAPA RNA HyperPrep Kit with RiboErase (HMR) (Roche Diagnostics). The adapter ligation step of library preparation involved a 1:10 dilution of NEBNext^®^ Multiplex Oligos for Illumina^®^ Sets 1, 2, and 3 (New England Biolabs) for multiplexing. For library amplification, a mixture composed of 3 *µ*L USER^®^ Enzyme (New England Biolabs) and 30 *µ*L of the library amplification master mix was combined with 17 *µ*L of the product from the second post-ligation cleanup, making a total reaction volume of 50 *µ*L. This mixture was then incubated at 37°C for 30 min prior to the thermocycling process. Following the completion of library preparation, the integrity and quality of the libraries were assessed using either the Bioanalyzer 2100 system (Agilent Technologies) or the TapeStation 2200 system (Agilent Technologies), and concentrations were determined via the Qubit fluorometer (ThermoFisher Scientific). The multiplexed library pools were quantified using the KAPA Library Quantification Kit (Roche Diagnostics) and subjected to sequencing. This sequencing was performed on the NovaSeq 6000 system (Illumina) using 150 bp paired end reads.

### 4.7 RNAseq data, gene expression analysis, and splicing variant analysis

RNA sequencing libraries were sent to Novogene for sequencing on the NovaSeq 6000 platform. Following sequencing, reads from technical replicates of the same sample were merged. The quality control process for these reads was performed using TrimGalore, trimming nucleotide bases with Phred scores below 20. The STAR aligner software (version 2.6.1d) was used to map the quality-controlled and trimmed reads against the human reference genome (GRCh38) using a 2-pass alignment mode. Here, the initial alignment pass maps the reads to the reference genome to identify splice junctions expressed in each sample. Subsequently, a second alignment pass is conducted, producing position-sorted binary alignment map (BAM) files. Based on the list of splice junctions detected across all samples in the previous round of alignment, the splice junction counts for each sample were generated.

For quantification of gene expression, BAM files were sorted based on read names. Gene read counts within these sorted BAM files were then determined using featureCounts. The quantification process included all genes listed in the GENCODE v31 gene transfer format (GTF) file. Subsequently, the raw gene counts were normalised to counts per million (CPM) to account for variations in library size across samples; this was achieved by adjusting the counts relative to the library size for each sample and then multiplied the result by 1,000,000. These normalised values were then employed as input data for differential gene expression analysis using DESeq2 (version 1.30.1). Significantly differentially expressed genes were identified if they have a log2 fold change of *> |*0.10*|*; expression in at least three samples within at least one of the two groups being compared in the differential expression analysis; and false discovery rate (FDR) below 0.01. Genes that were not expressed at sufficient levels, defined as having a CPM value of less than 1, were excluded from the analysis. Finally, enriched biological pathways among the differentially expressed genes were identified using ClusterProfiler (version 3.18.1).

To analyse splicing variants, StringTie (version 2.1.4) was used to assemble a GTF file listing splicing variants identified across all samples examined. Individual sample GTF files were then merged into a comprehensive GTF file containing all splicing variants identified in this study. Novel splicing variants not previously documented in the GENCODE v31 database were identified using tailor-made R scripts. For the detection of alternative splicing at the exon junction level, rMATS (version 4.1.0) was used to identify novel splicing events, based on the known and newly identified splicing variants. Within this framework, rMATS categorized alternative splicing events into five types: skipped exon (SE), retained intron (RI), mutually exclusive exons (MXE), alternative 5’ splice sites (A5SS), and alternative 3’ splice sites (A3SS).

Following the identification of these alternative splicing categories, MARVEL (Modality Assessment to ReVEaL alternative splicing dynamics at single-cell resolution), an R package designed for alternative splicing analysis in both bulk and single-cell samples, was used for differential splicing analysis. This analysis takes splice junction read counts (from STAR aligner), alternative splicing events counter (from rMATS), and the GENCODE v31 GTF catalogue, to compute a percent spliced-in (PSI) value for each splice junction, reflecting the proportion of reads indicative of alternative splicing relative to the total reads at that junction.

Differential splicing analysis was then conducted on adequately expressed splicing events, defined as those supported by a minimum of 10 reads across various splicing variants. For each event, the difference in permuted PSI values (ΔPSIperm) was calculated 1,000 times to establish a null distribution. The absolute percentage of values within the null distribution that is larger than the observed ΔPSIobs were recorded and represents the *P* value of the ΔPSIobs. Splicing events were considered significantly differentially spliced if they met two criteria: a P-value *<* 0.01 and an absolute ΔPSIobs greater than *|*0.15*|*.

### 4.8 Hybridisation chain reactions (HCR) staining, imaging, and quantification

On the day of HCR staining, samples were washed twice with 300 *µ*L of 2× SSC. Samples were pre-hybridised in 300 *µ*L of probe hybridization buffer for 30 min at 37°C. Probe solution was prepared by adding 1.2 pmol of each probe set (e.g. 1.2 *µ*L of 1 *µ*M stock) to 300 *µ*L of probe hybridization buffer at 37°C, then pre-hybridization solution was replaced with probe solution. Samples were incubated overnight (12–16 h) at 37°C. The next day, excess probe was removed by washing 4 × 5 min with 300 *µ*L of probe wash buffer at 37°C. Samples were washed 2 × 5 min with 5× SSCT at room temperature. Samples were pre-amplified in 300 *µ*L of amplification buffer for 30 min at room temperature. 18 pmol of hairpin h1 and 18 pmol of hairpin h2 were separately prepared by snap cooling 6 *µ*L of 3 *µ*M stock (heat at 95°C for 90 seconds and cool to room temperature in a dark drawer for 30 min). Hairpin solution was prepared by adding snap-cooled h1 hairpins and snap-cooled h2 hairpins to 300 *µ*L of amplification buffer at room temperature, then pre-amplification solution replaced the hairpin solution. Samples were incubated overnight (12–16 h) in the dark at room temperature. Excess hairpins were removed by washing 5 × 5 min with 300 *µ*L of 5× SSCT at room temperature. Then SSCT was replaced with *∼*100 *µ*L of antifade mounting reagent. Samples were stored at 4°C protected from light prior to imaging. Images were acquired with a Zeiss LSM-980 laser scanning confocal microscope. This microscope is equipped with an inverted Axio Observer microscope stand, 2 multialkali PMT detectors, and 32 GaAsP PMT detectors. Fluorescent labels were excited with either 405 nm, 488 nm, 561 nm or 639 nm diode lasers. The microscope was configured with a Zeiss C-Apochromat 40x/1.2 W Corr objective for imaging. HCR signal was validated using the 32 GaAsP PMT detectors in spectral imaging mode covering wavelengths ranging from 411 nm to 694 nm with a resolution of 8.8 nm. Images were processed in Fiji using a rolling ball background subtraction (radius = 50 pixels) followed by a gaussian blur filter (sigma (radius) = 0.8).

### 4.9 Synthesis of CRISPR-Lock AAV, transgenic mice, and AAV injections

Ultra-purified AAV9 vectors encoding either the full-length dPspCas13b along with its 23 nt repeat-targeting crRNA (construct 1), the compact CRISPR-Lock 2, or control AAV were prepared by VectorBuilder. HSA^LR^ transgenic mice harboring more than 200 CTG repeats were selected for experimental procedures. The HSA^LR^ mice were housed and bred at the Department of Biomedical Services, University of Oxford. Mice were housed in a light/dark cycle with access to food and water ad libitum. The mice were subjected to inhalation anesthesia, initiated with 3% isoflurane and maintained at 1.5% isoflurane throughout the procedure. For the tibialis anterior (TA) muscle injections, AAV vectors were thawed on ice, diluted in sterile phosphate-buffered saline (PBS), and then administered into the TA muscles of the anesthetized HSA^LR^ mice, which were between 5 and 9 weeks old. A 29-gauge needle was used to deliver the AAV across three distinct injection sites within the TA muscles.

### 4.10 Mouse tissue collection, total RNA isolation, and ddPCR assay

After AAV intramuscular injection, TA muscles were harvested from treated mice and processed by homogenization in a 1-thioglycerol/homogenization solution using a Precellys tissue homogenizer (Bertin Instruments) for two intervals of 1 minute each. Following this, total RNA was isolated with the Maxwell RSC simplyRNA Tissue Kit (Promega) using the Maxwell RSC Instrument (Promega), according to manufacturer’s instructions. Total RNAs were isolated using the PureLink RNA Mini Kit (Life Technologies) following the manufacturer’s instructions. Purified total RNAs were then subjected to DNase treatment using the TURBO DNA-free™ Kit (ThermoFisher Scientific) in a total volume of 30 *µ*L, as per the manufacturer’s protocol. cDNA was synthesized from 75 ng of the treated RNA for each sample using the SuperScript™ First-Strand Synthesis System (Life Technologies), which was then used in ddPCR analyses.

The primers and probes for the ddPCR assays were custom-designed and synthesized by Integrated DNA Technologies (IDT), with the sequences detailed in Supplementary Table 3. ddPCR reaction mixtures were prepared using the ddPCR™ Supermix for Probes (BioRad), according to manufacturer’s instruction. Each batch of ddPCR included a no-template control to monitor for any potential contamination. The ddPCR mixture was then loaded into a DG32 cartridge and droplets were formed using the Automated Droplet Generator (BioRad). Carefully, 40 *µ*L of the droplet-oil emulsion was transferred to a 96-well plate, which was then heat-sealed with foil. The sealed plate was subjected to PCR amplification in a C1000 Touch Thermal Cycler (BioRad), following the specified cycling con-ditions in the ddPCR™ Supermix for Probes kit. Following PCR, the droplets in each sample were quantified using a QX200 droplet reader, and data were analysed with QuantaSoft software to calculate the count of positive droplets. Serial dilutions of cDNA were performed in each ddPCR assay to achieve a minimal count of double-positive droplets. Additionally, for each primer pair, gradients of annealing temperatures were tested to optimise the PCR conditions and maximise the dynamic range of the assay.

### 4.11 FISH-immunofluorescent staining of muscle cryosections

TA muscle cryosections, with thicknesses ranging from 8 to 10 *µ*m, were fixed using 4% PFA in PBS for 10 minutes. Muscle sections were then washed three times with PBS before being permeabilized with 0.5% Triton in PBS at room temperature for 10 min. After removing the Triton solution, sections were rehydrated in a wash buffer composed of 50% formamide and 2× SSC buffer for 10 minutes. Subsequently, sections were treated with a hybridization buffer containing 5% dextran sulfate, 2× SSC buffer, 0.02% BSA, 1 *µ*g/*µ*l yeast tRNA, formamide (at half the final volume), and water to achieve the desired volume for 15 minutes at 37°C in a hybridization oven. A Cy3-labeled 2*^′^*OMe (CAG)_7_ probe (IDT) was denatured at 98°C for 10 minutes, and 0.2*µ*M of this probe was mixed into the pre-chilled hybridization buffer before being immediately added to the sections. Hybridization was carried out for 2 h at 37°C in a hybridization oven. Following hybridization, sections were washed three times with wash buffer for 30 min each, then rinsed once with PBS, and finally, slides were mounted using a mounting medium containing DAPI. For experiments combining fluorescence in situ hybridization (FISH) with immunofluorescence, immunostaining was conducted after the final FISH wash. Immunostaining used the rabbit polyclonal anti-MBNL1 antibody (kindly provided by C. Thornton; diluted 1:2000) and a secondary 1:500 Alexa Fluor 647-conjugated goat anti-rabbit antibody (Life Technologies). The distribution of FISH foci was assessed using a confocal microscope (FV1000), and the automated quantification of nuclei containing foci, the number of foci per nucleus, and the intensity of these foci was done with Imaris image analysis software (Oxford Instruments).

### 4.12 Statistical analyses

Unless otherwise stated, all assays were performed in at least three independent biological replicates. Two independent biological replicates were performed in the evaluation of single plasmid format for CRISPR-Lock. In the mouse experiments, the statistical significance was determined using one-way ANOVA followed by Dunnett’s test. All the *P* values are shown for each of the groups in comparison with the negative control group. All calculation and statistical analyses of the experimental and computational data were conducted using GraphPad Prism v.10.0.2 and R, respectively. Statistical significance between different groups was calculated using the tests indicated in each figure legend. No statistical methods were used to predetermine sample size.

## 5 Acknowledgements

We thank the WIMM Flow Cytometry facilities (Paul Sopp, Sally-Ann Clark, Craig Waugh, Kevin Clark), the MRC WIMM Centre for Computational Biology and WIMM Wolfson Imaging Centre, the Dunn School Imaging facilities and Oxford Biomedical Services. We also thank Sauka-Spengler lab (Andrew Ramos, Oana Pelea, Sarah Mayes, Mara Artibani, Adi Harlianto, Jasmina Kuburic) and Desiree Wilson for technical advice and support, and colleagues at the lab and the WIMM for academic discussion (Tom Milne, David Knapp, Quentin Ferry, Yale Michaels, Daniel Fountain, and Evgeny Akkuratov). We are grateful to Sabbi Lall for advice on the manuscript.

## 6 Funding

This work was supported by the Wellcome Trust Grant (ref 215615/Z/19/Z) and the institutional support from Stowers Institute for Medical Research to T.S.S.. M.H. was supported by the Boehringer Ingelheim Fonds Fellowship.

## 7 Author Contributions

M.H., T.S.S., and T.A.F. conceptualised the project. M.H., T.S.S., and T.A.F. developed the methodology. M.H., P.S.A.K., M.F., C.R. performed the experiments. M.H., P.S.A.K., S.W., A.K., Z.Y., S.T., A.M., analysed data and generated data visualisation. P.S.A.K., Z.Y., A.K., performed HCR imaging experiments. S.W. performed computational analysis of RNA-seq data. M.F. performed mouse injections, and M.W. provided access to the DM1 mouse model. M.H. and T.S.S. acquired funding. T.S.S. administered the project. T.S.S. and C.R. supervised the project. M.H. wrote the original draft. M.H. and T.S.S. reviewed and edited the draft.

## Extended Data

**Extended Data Figure 1:**
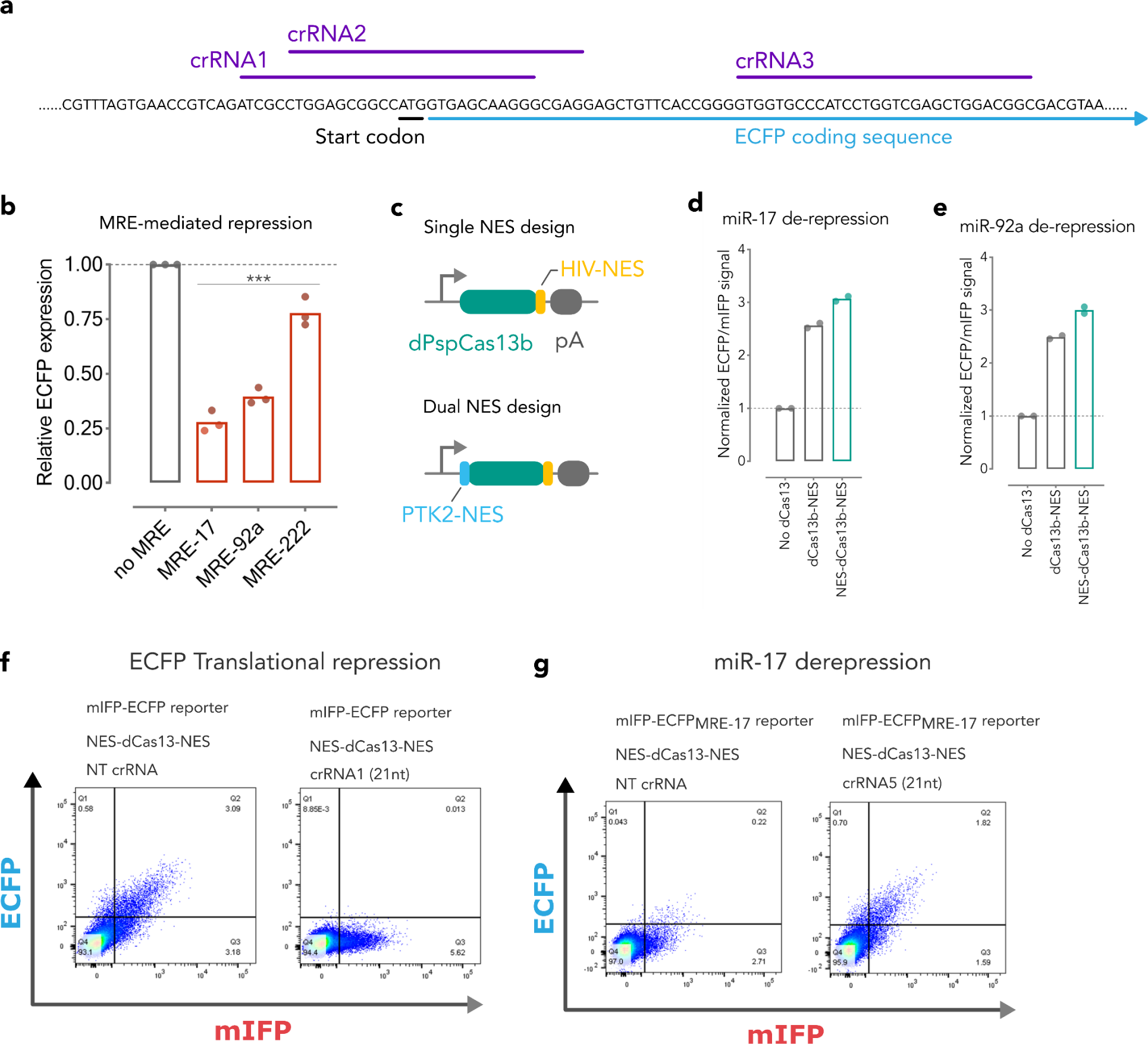
Design rule to improve specificity of CRISPR-Lock 1 for perturba-tion of individual miRNA-target interactions. **a**, Schematic diagram depicting a strategy to increase the specificity of CRISPR-Lock systems for targeting individual miRNA-target interactions. miRNAs typi-cally work in the context of complex regulatory networks, where each miRNA species has hundreds of MREs, presenting a challenge for precise perturbation of individual miRNA-target interactions with CRISPR-Lock. Designing crRNA (green) that partially covers the MRE (red) as well as its unique neighbouring region (shades of grey) enables perturbation of individual miRNA-target interactions. **b**, Schematic of experimen-tal design to evaluate mismatch tolerance profile of CRISPR-Lock system. crRNAs are designed to pair with the MRE and the MRE-neighbouring region of the target transcript. Using these designs, evaluation of CRISPR-Lock 1 steric blocking efficiencies in target and non-target transcripts (same MRE sequence, but different MRE-neighbouring sequence) was performed. **c**, Design of crRNAs that are complementary to the unique MRE-neighbouring region either in their 5’ side (p10 – p12) or 3’ side (p13 – p15). Dot-ted green lines depict spacer regions covering MRE-neighbouring regions. Green lines depict spacer regions covering the MRE (red). Orange shade depicts miRNA seed-matching region of the MRE. **d**, Schematic of the CRISPR-Lock 1 crRNAs as indicated in **c**, highlighting the position and extent of spacer mismatches in each of the designed crRNA (p10 – p15). **e-h**, Bar graphs showing the efficiency of CRISPR-Lock 1 in de-repressing miRNA mediated silencing in either target RNAs (green) or non-target RNAs (red) (n = 3).

**Extended Data Figure 2:**
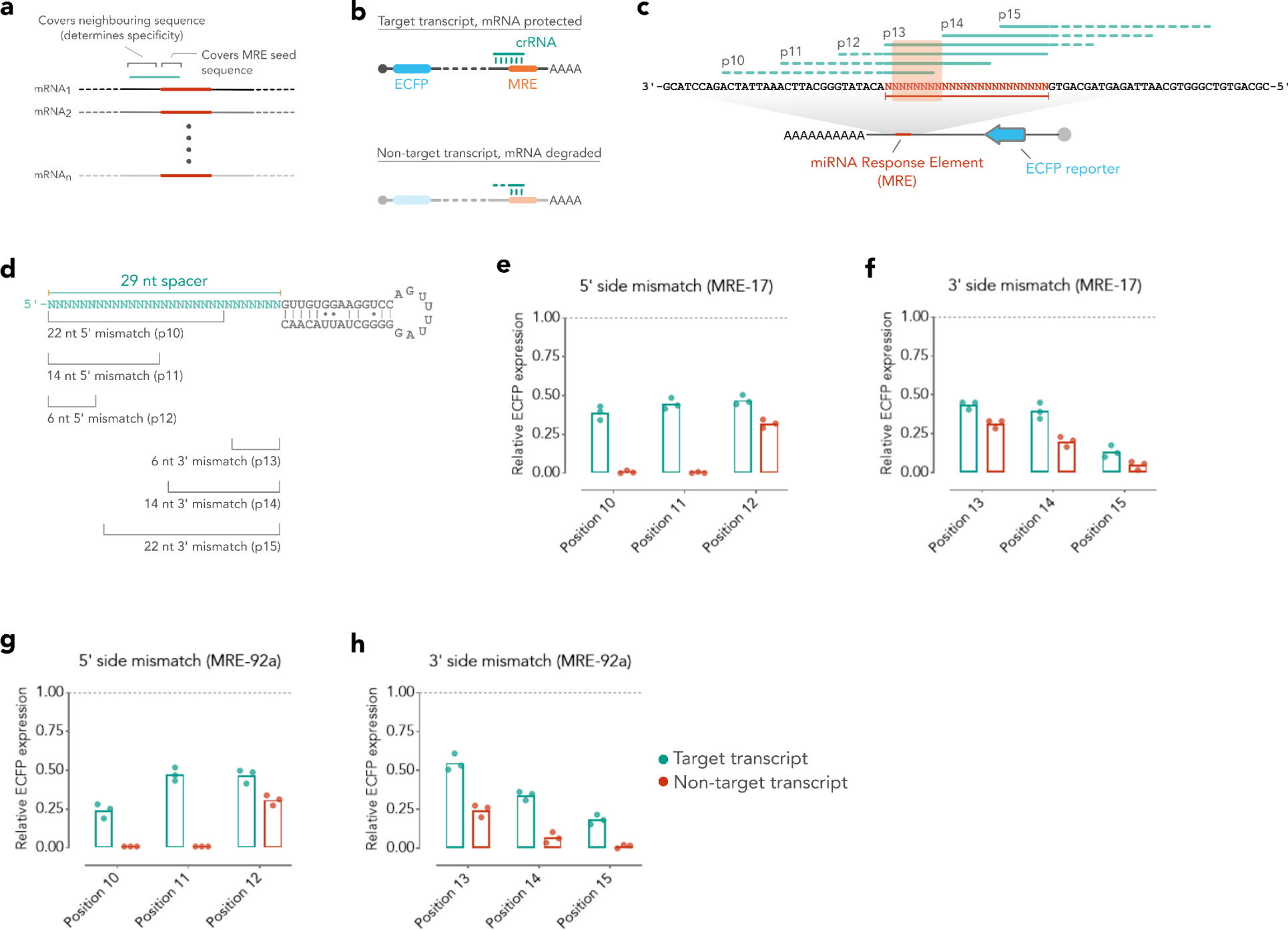
Design rule to improve specificity of CRISPR-Lock 1 for perturbation of individual miRNA-target interactions. **a**, Schematic diagram depicting a strategy to increase the specificity of CRISPR-Lock systems for targeting individual miRNA-target interactions. miRNAs typically work in the context of complex regulatory networks, where each miRNA species has hundreds of MREs, presenting a challenge for precise perturbation of individual miRNA-target interactions with CRISPR-Lock. Designing crRNA (green) that partially covers the MRE (red) as well as its unique neighbouring region (shades of grey) enables perturbation of individual miRNA-target interactions. **b**, Schematic of experimental design to evaluate mismatch tolerance profile of CRISPR-Lock system. crRNAs are designed to pair with the MRE and the MRE-neighbouring region of the target transcript. Using these designs, evaluation of CRISPR-Lock 1 steric blocking efficiencies in target and non-target transcripts (same MRE sequence, but different MRE-neighbouring sequence) was performed. **c**, Design of crRNAs that are complementary to the unique MRE-neighbouring region either in their 5’ side (p10 – p12) or 3’ side (p13 – p15). Dotted green lines depict spacer regions covering MRE-neighbouring regions. Green lines depict spacer regions covering the MRE (red). Orange shade depicts miRNA seed-matching region of the MRE. **d**, Schematic of the CRISPR-Lock 1 crRNAs as indicated in **c**, highlighting the position and extent of spacer mismatches in each of the designed crRNA (p10 – p15). **e-h**, Bar graphs showing the efficiency of CRISPR-Lock 1 in de-repressing miRNA mediated silencing in either target RNAs (green) or non-target RNAs (red) (n = 3).

**Extended Data Figure 3:**
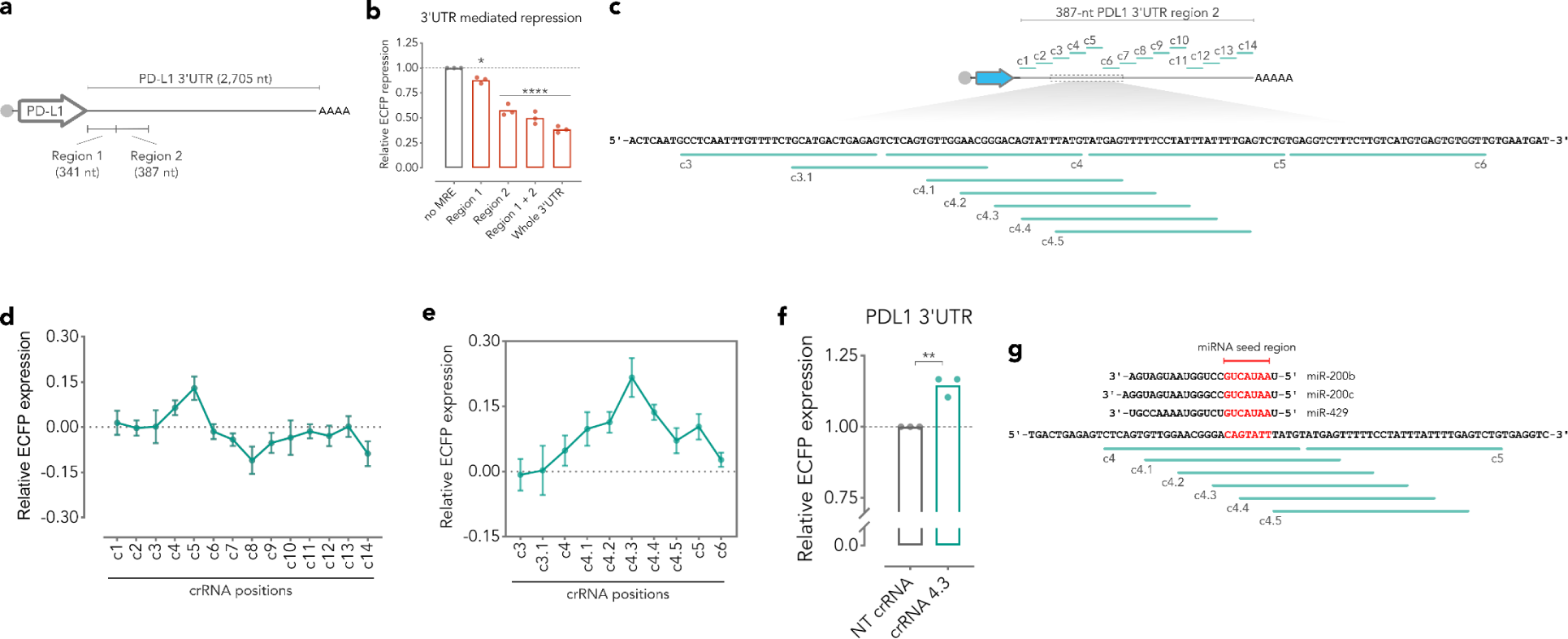
Proof of principle application of CRISPR-Lock 1 for screening RNA functional elements. **a**, Schematic diagram depicting PD-L1 3’UTR. **b**, Bar graphs showing the silencing of ECFP reporter containing either full or partial sequence of PD-L1 3’UTR. Strong repression mediated by Region-2 indicates the presence of a strong de-stabilizing element. **c**, Schematic diagram of the CRISPR-Lock 3’UTR screening strategy in Region-2 reporter to identify de-stabilizing elements in PD-L1 3’UTR. **d**, Line graph showing upregulation or downregulation of ECFP expression in CRISPR-Lock screening experiment. Functional destabilizing elements were predicted to be present in the 3’UTR fragment covered between crRNA-3 to crRNA-6. crRNAs used in this experiment are designed in a non-overlapping manner. **e**, Line graph showing upregulation of ECFP expression from fine-resolution CRISPR-Lock screening experiment, where crRNAs are designed in 5 nucleotide shifts pattern (crRNA 4.1 to crRNA 4.5). The strongest ECFP upregulation was observed in steric inhibition mediated by crRNA 4.3 guided CRISPR-Lock 1. **f**, Upregulation of ECFP reporter expression upon delivery of CRISPR-Lock 1 components guided by crRNA-4.3 (c4.3) compared to NT control. **g**, Cross-examination of PD-L1 3’UTR sequence covered between crRNA 4 and crRNA 5 reveals the presence of miRNA seed-complementary sequence (red).

**Extended Data Figure 4:**
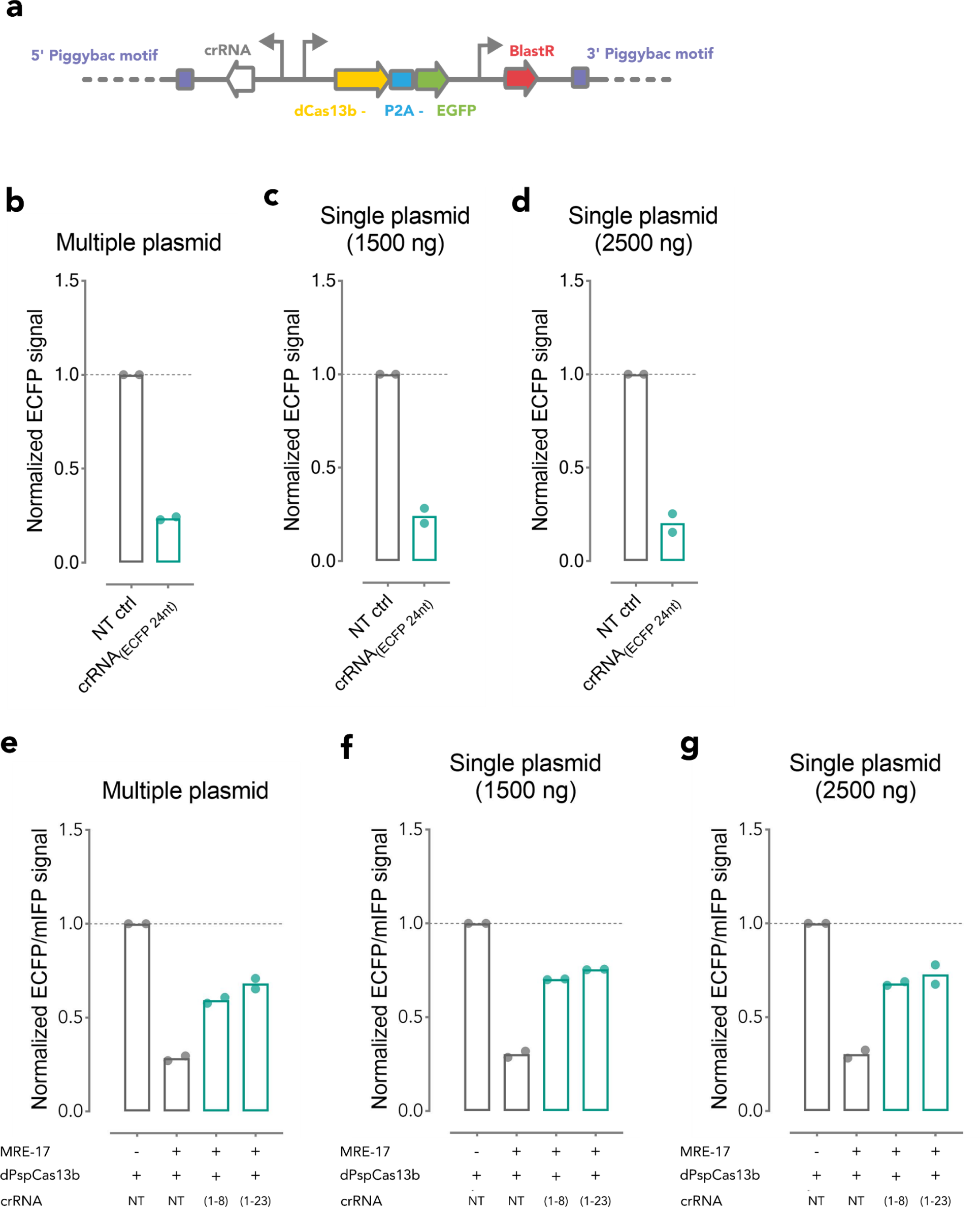
Design and evaluation of single plasmid format for CRISPR-Lock. **a**, Schematic diagram showing construction of single plasmid CRISPR-Lock. Human U6 promoter (pU6) drives the expression of crRNA in an opposite orientation to EF1*α* promoter (pEF1*α*)-driven dPspCas13. **b-d**, Steric blocking efficiency of CRISPR-Lock 2 targeting Kozak sequence of the ECFP reporter in multiple-plasmids (500 ng dPspCas13-encoding and 1,000 ng crRNA-encoding plasmids) or single-plasmid format. **e-g**, Steric blocking efficiency of CRISPR-Lock 1 targeting MRE-17 silencing the expression of the ECFP reporter in multiple plasmids (500 ng dPspCas13-encoding and 1,000 ng crRNA encoding plasmids) or single-plasmid format. CRISPR-Lock systems encoded in a single-plasmid format resulted in similar steric blocking efficiency compared to multiple-plasmids format.

**Extended Data Figure 5:**
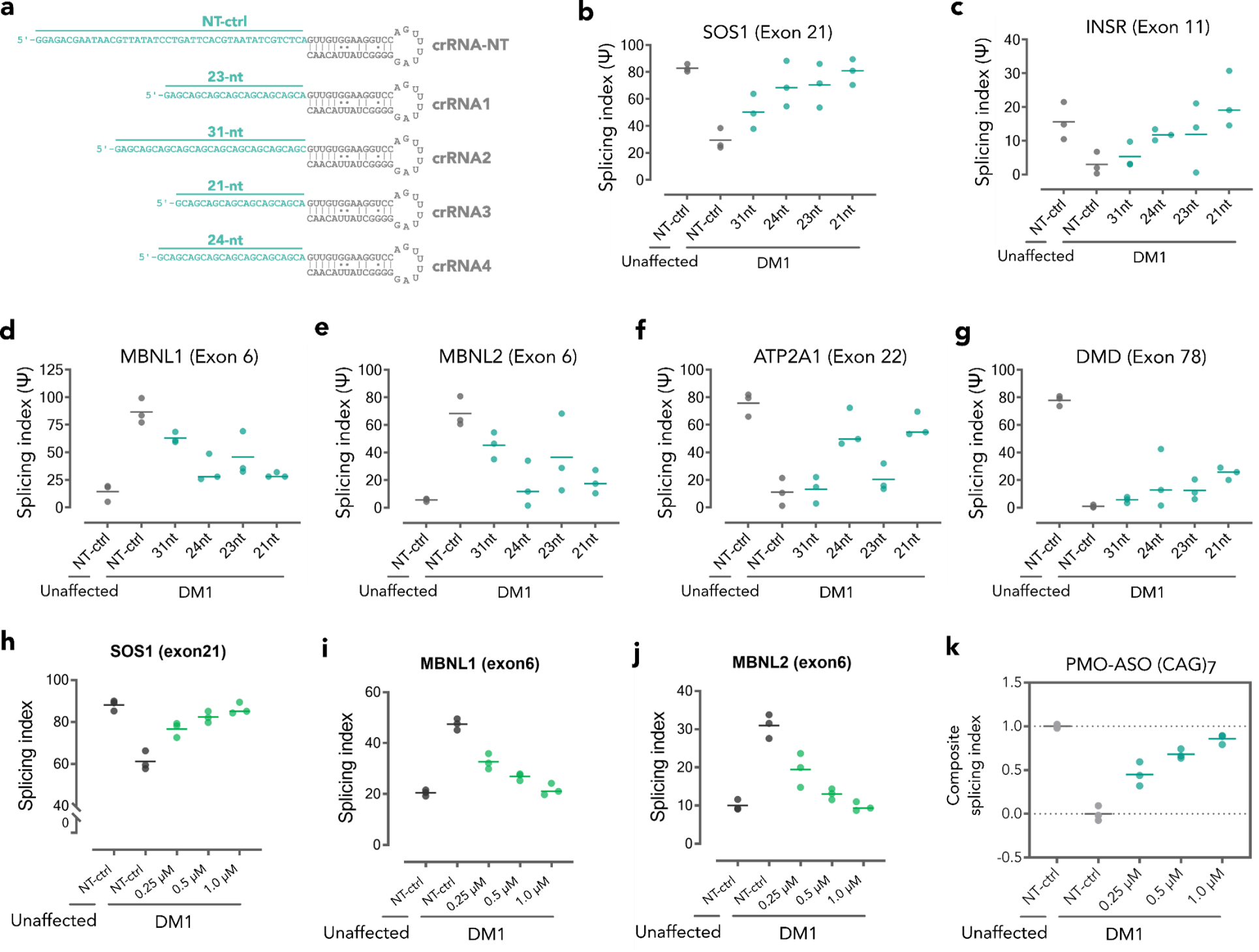
Correction of DM1-associated mis-splicing biomarkers after treatment with (CUG)n-targeting PMO-ASO or CRISPR-Lock 2. **a**, Schematic diagram of five crRNAs used in this experiment. **b-g**, Steric inhibition of (CUG)n repeat expanded sequence in DM1 patient-derived cells resulted in correction of splicing index in 6 clinically relevant DM1 biomarkers, which are SOS1 (**b**), INSR (**c**), MBNL1 (**d**), MBNL2 (**e**), ATP2A1 (**f**), and DMD (**g**). Treatment with CRISPR-Lock 2 guided by crRNAs containing short 21-nt spacer resulted in highest levels of corrections. **h-j**, Intracellular delivery of (CAG)7 PMO-ASO into patient-derived cells resulted in dose-dependent correction of clinically relevant splicing dysfunction biomarkers, which are SOS1 (**h**), MBNL1 (**i**), and MBNL2 (**j**). Intracellular delivery of 1 µM (CAG)7 PMO-ASO led to complete correction of these biomarkers (**k**). In **h-j**, and **b-g**, Plotted values are quantification of the band intensity from semiquantitative PCR assay from n = 3 independent replicates.

**Extended Data Figure 6:**
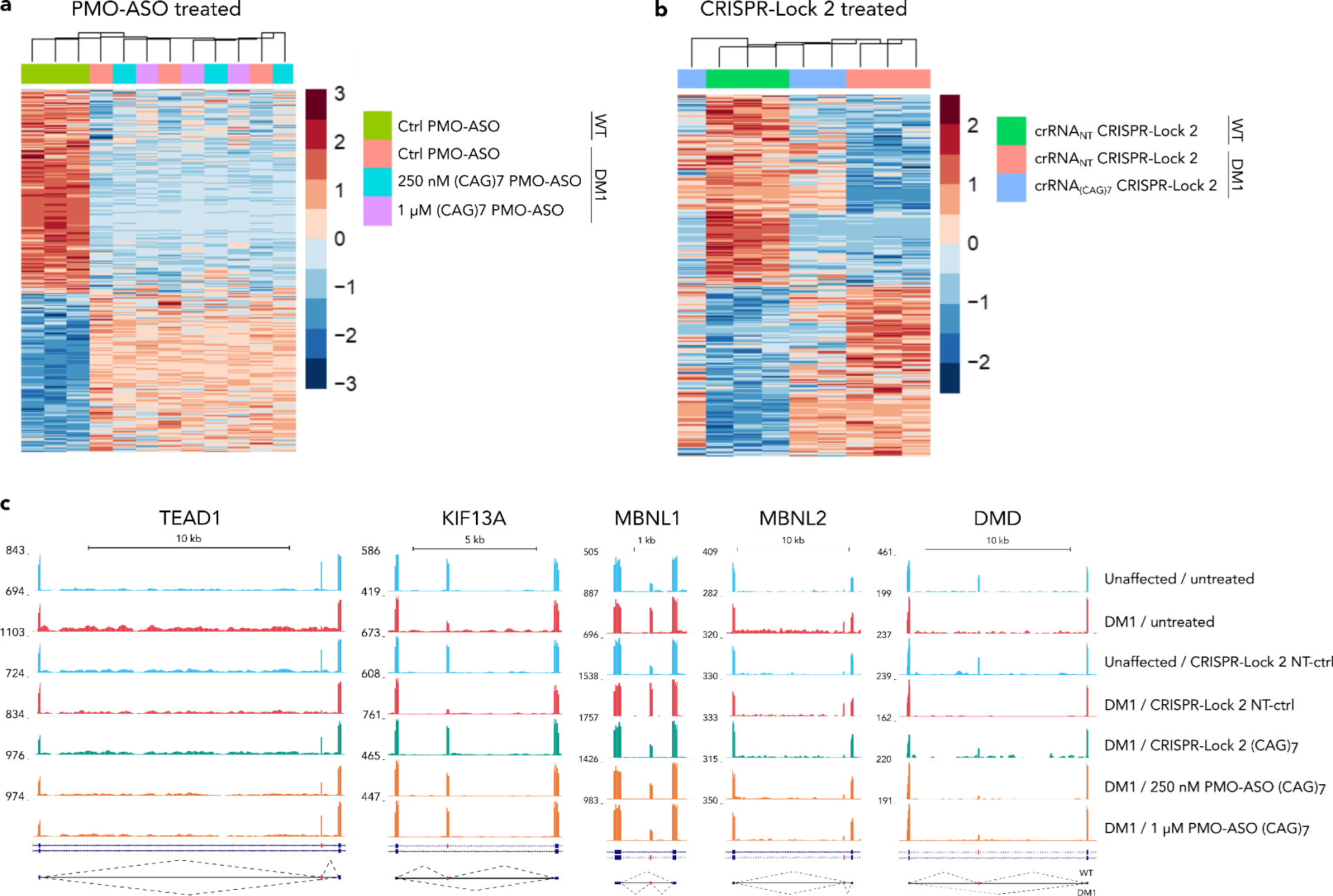
CRISPR-Lock 2 reverses mis-splicing on key transcription factors, splicing factors, and a structural protein in DM1 cells. **a**, Hierarchical clustered heatmap of splicing indices of high-confidence splicing dysregulation events in WT (unaffected) and DM1 myotubes treated with (CAG)7 PMO-ASO or control PMO-ASO. **b**, Hierarchical clustered heatmap of splicing indices of high-confidence splicing dysregulation events in WT (unaffected) and DM1 myotubes treated with CRISPR-Lock 2 guided with either crRNA(CAG)7 or NT-crRNA. **c**, RNA-seq data represented on UCSC genome-browser tracks showing correction of DM1-related splicing dysregulation events in TEAD1, KIF13A, MBNL1, MBNL2, and DMD transcripts in myotubes from WT or DM1 cells treated with either CRISPR-Lock 2 or PMO-ASO, as indicated.

**Extended Data Figure 7:**
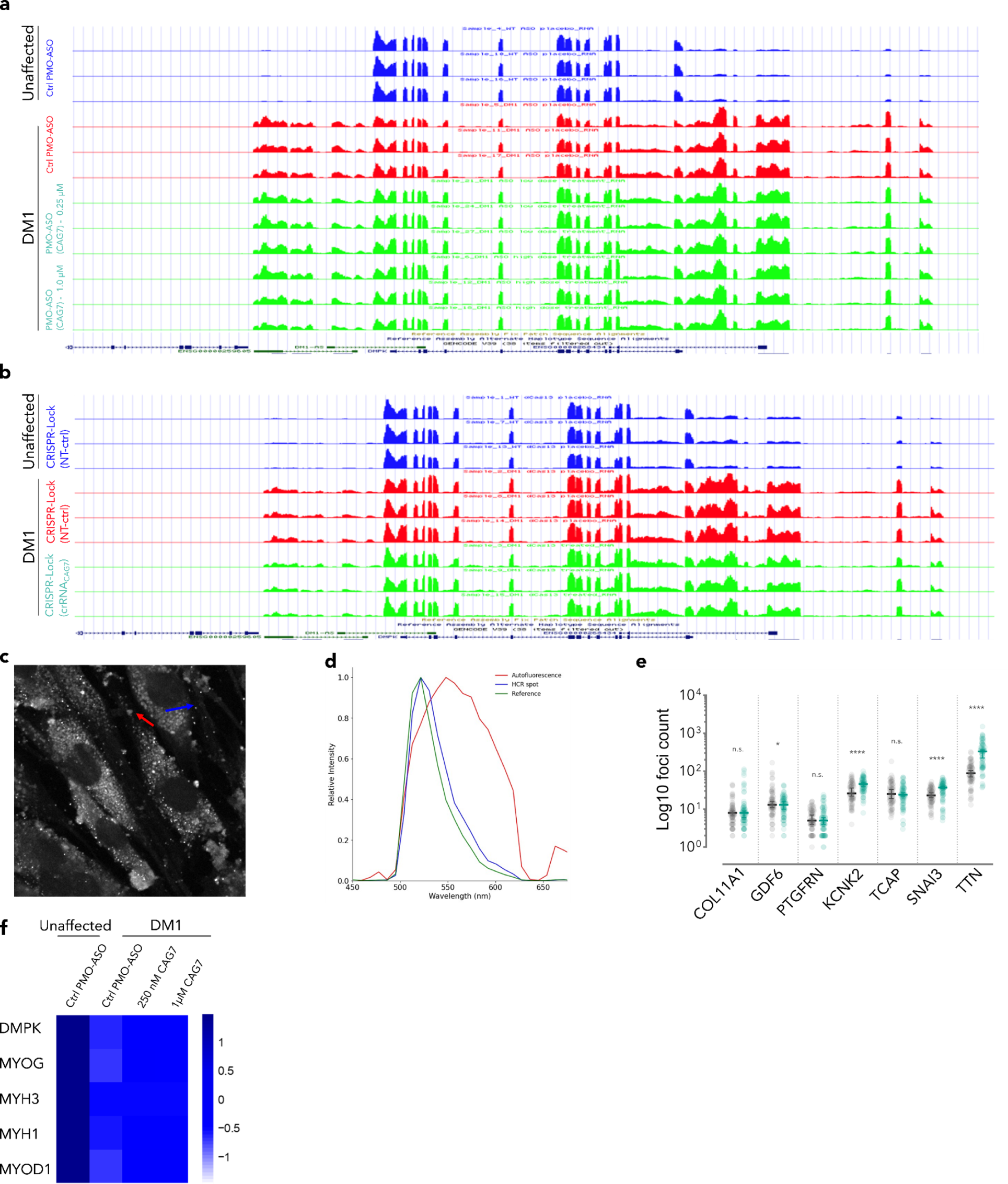
Expression level of DMPK transcript in PMO-ASO or CRISPR-Lock 2 treated myotubes and HCR image analysis and quantification. **a**, RNA-seq data represented on UCSC genome-browser tracks showing the expression level of DMPK transcripts in unaffected and DM1 cells treated with PMO-ASO. **b**, RNA-seq data represented on UCSC genome-browser tracks showing the expression level of DMPK transcripts in unaffected and DM1 cells treated with CRISPR-Lock 2. **c-d**, Discriminating HCR spot signal from background auto fluorescence. **c**, Visual discrimination of HCR signal (blue arrow) and background auto fluorescent (red arrow). **d**, Measurement of wavelength and signal intensity of HCR signal and background auto fluorescent from (**c**). **e**, Quantification of HCR signals from structural proteins and transcription factors associated with muscle tissue differentiation in myotubes treated with CRISPR-Lock 2 guided with either non-targeting crRNA (grey) or crRNA(CAG)7 (green). **f**, Expression level of DMPK transcript and four muscle differentiation markers in human myotubes treated with either ctrl PMO-ASO or (CAG)7 PMO-ASO, as evaluated by RNAseq. NT, non-targeting; PMO-ASO, Phosphorodiamidate morpholino – antisense oligonucleotide; (CAG)7, 21-nt sequence targeting (CUG)n repeat expansion. In (**e**) *P* values are by two-tailed Student’s t-test. \**P <* 0.05; \*\**P <* 0.01; \*\*\**P <* 0.001; \*\*\*\**P <* 0.0001; n.s., not significant.

**Extended Data Figure 8:**
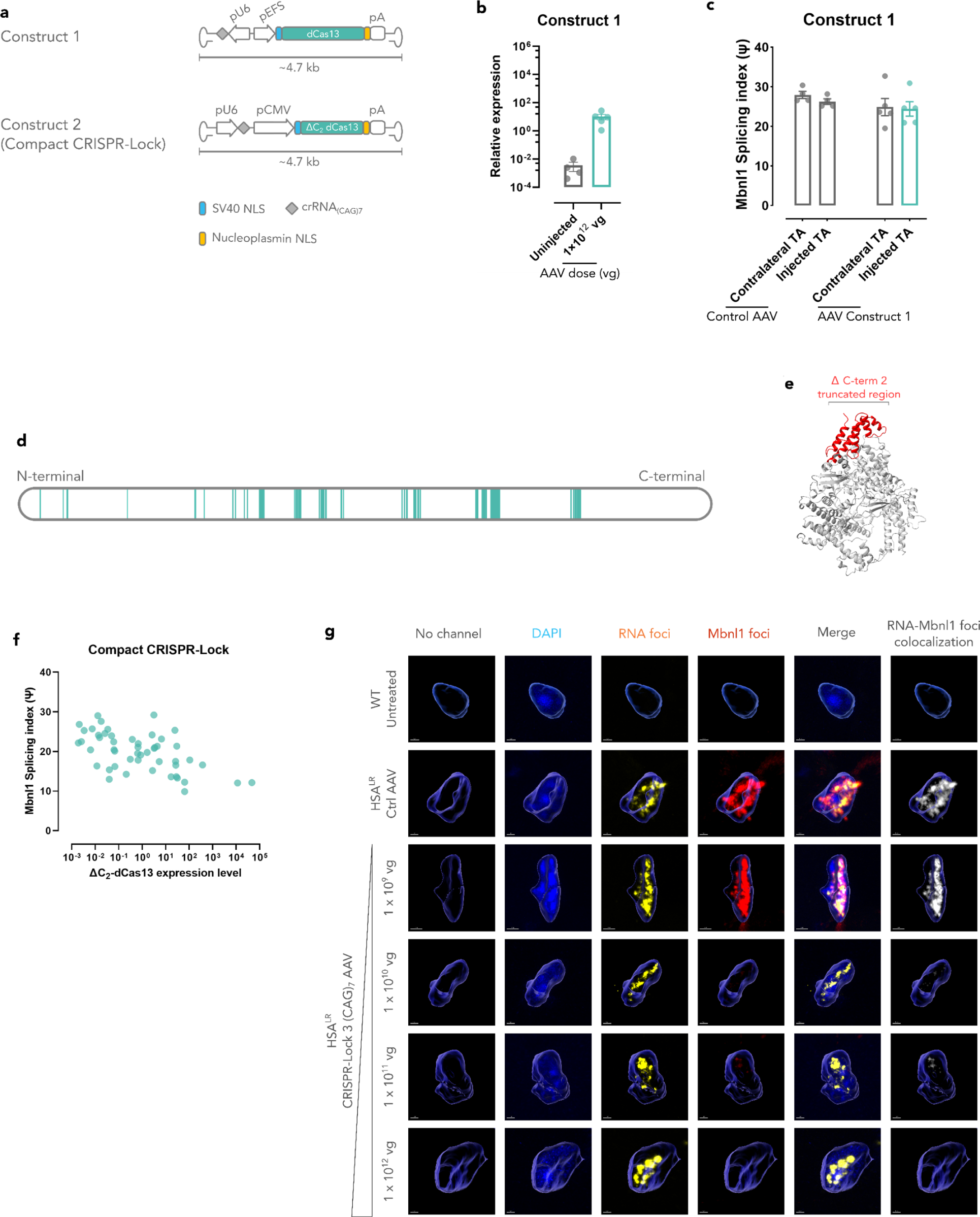
Predicted structure and crRNA-interacting amino acids and characterization of full-length CRISPR-Lock 2 AAV construct. **a**, Diagram showing the AAV-encoded pEFS-driven construct 1 (top) and pCMV-driven compact CRISPR-Lock. **b**, Bar graph showing the *in vivo* expression level of dPspCas13b in HSA^LR^ mice upon IM injection of construct 1 AAV. **c**, Bar graph showing the extent of *in vivo* correction of Mbnl1 splicing index upon injection of either control AAV or Construct 1 AAV. (control AAV group n = 4; Construct 1 AAV group n = 5). **d**, Diagram highlighting the location of the predicted crRNA-binding amino acids in dPspCas13b obtained from the alignment of dPspCas13b amino acid sequence to the database of conserved protein domain in Cas13b family. **e**, Prediction of dPspCas13b structure with AlphaFold 2, highlighting the truncated HEPN2 domain (red) in the compact CRISPR-Lock 2 construct. **f**, Dot plot showing the relationship between the expression level of Δ_2_-dCas13 with the correction of splicing dysregulation in Mbnl1 (n = 52). **g**, Representative nucleus showing dose dependent elimination of DM1-associated foci in HSA^LR^ muscle tissue after treatment with single-AAV CRISPR-Lock 2. Repeat-expanded (CUG)n foci are labelled with FISH (Cy3 channel), Mbnl1 foci were labelled with immunofluorescent (AF647 channel), nuclei are labelled with DAPI.

## Notes

### Competing Interest Statement

M.H. and T.S.S are co-founders of Eterna Biologics. C.R. is a co-founder of Isogenix. M.J.A.W is a co-founder of Isogenix.

