## Supplementary Information for "Robust CRISPR/dCas13 RNA blockers specifically perturb miRNA-target interactions and rescue type 1 myotonic dystrophy pathology"

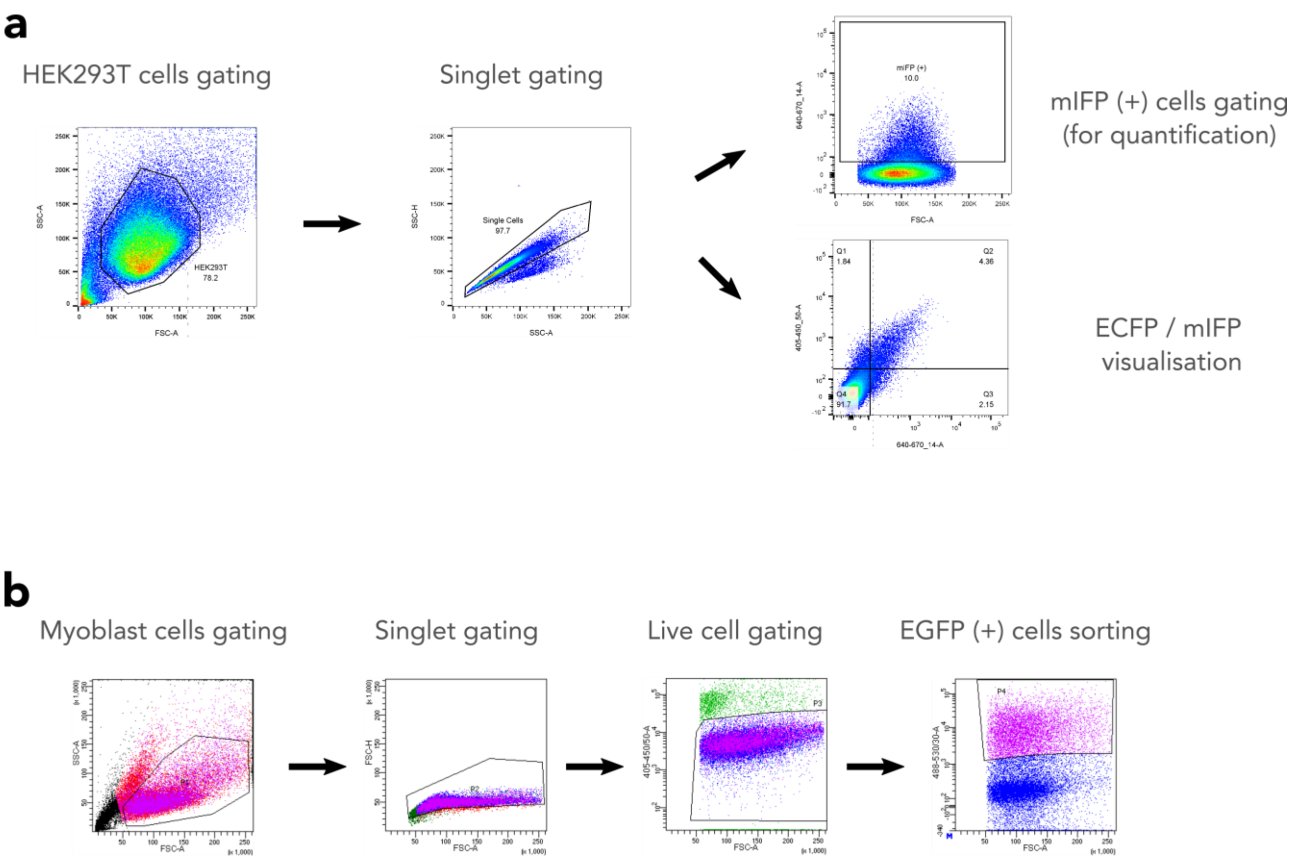


**Supplementary Figure 1. Representative gating strategies in translational repression, miRNA derepression, and DM1 treatment experiments. a**, gating hierarchy used in all bidirectional reporter experiments, including CRISPR-Lock translational repression and miRNA derepression experiments. In the first gate, HEK293T population was selected based on their FSC-A (x-axis) and SSC-A (y-axis) values. In the second gate, singlets were selected based on their SSC-A (x-axis) and SSC-H (y-axis) values. In the final gate, a population of transfected cells were selected for the quantification of reporter expression in translational and miRNA derepression experiments. Alternatively, to generate representative dot plots to visualise CRISPR-Lock blocking efficiency, singlets were plotted based on expression level of ECFP (y-axis) and mIFP (x-axis). **b**, gating hierarchy used in experiments involving treatments of WT and DM1 myoblasts with CRISPR-Lock systems. In the first gate, myoblast population was selected based on their FSC-A (x-axis) and SSC-A (y-axis) values. In the second gate, singlets were selected based on their FSC-A (x-axis) and FSC-H (y-axis) values. In the third gate, live cells were selected based on the absence of DAPI staining in this cell population. In the final gate, transfected myoblasts were sorted based on the expression level of dCas13-2A-EGFP.

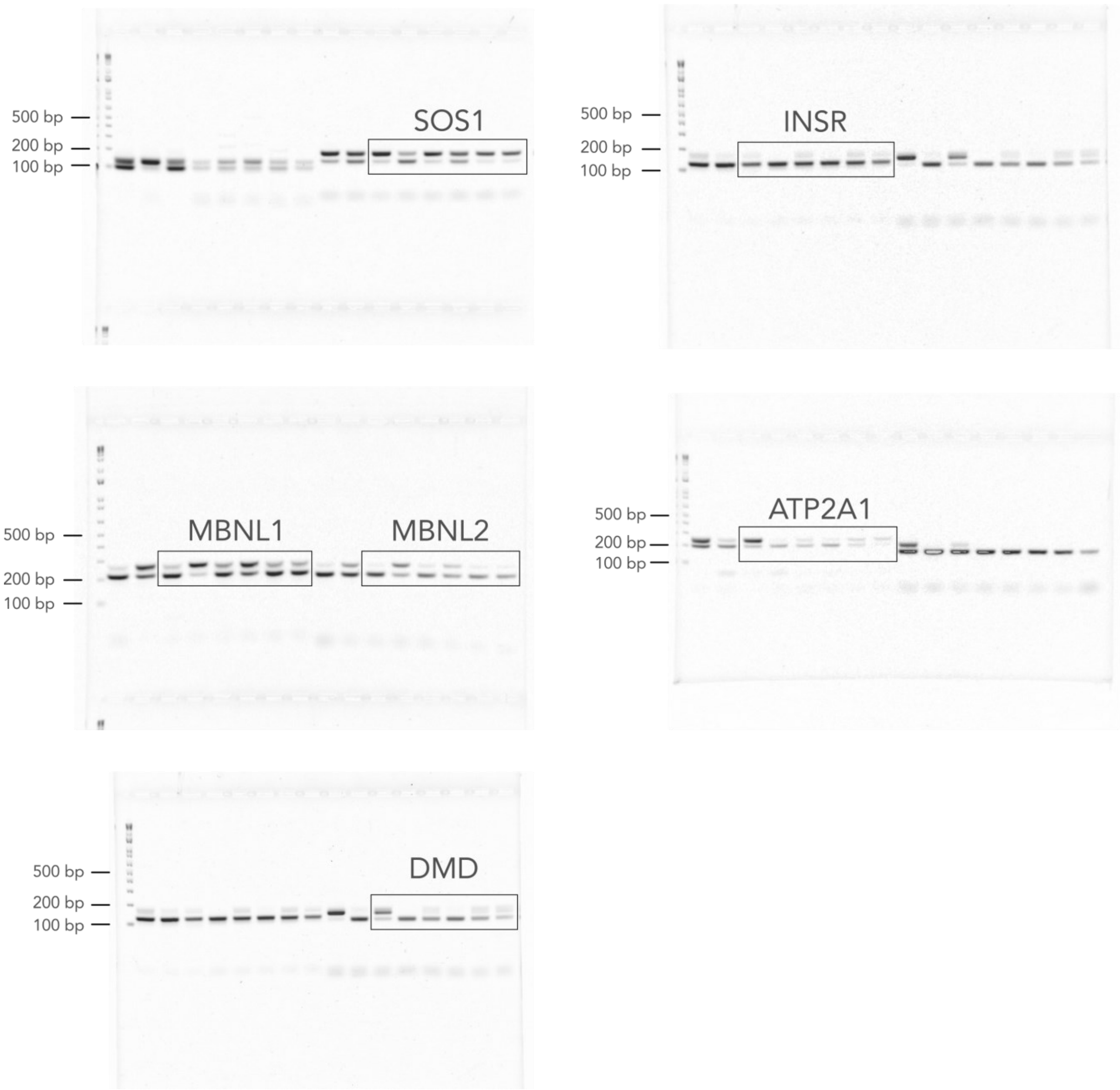


**Supplementary Fig. 2. Uncropped original gel electrophoresis data.** 1kb plus DNA ladder was used as a standard, labelled as 100 bp, 200 bp, and 500 bp. Gels were used to generate Fig 3c, with a black outline to show the boundary of the cropped portion.
